# Proximity biotinylation at the host-*Shigella* interface reveals UFMylation as an antibacterial pathway

**DOI:** 10.1101/2025.05.29.656827

**Authors:** Ana T. López-Jiménez, Fabien Théry, Kathryn Wright, Hannah Painter, Shelby T. Hoffmeister, Lucas Jarche, Jeremy Benjamin, Gerbrand J. van der Heden van Noort, Dominik Brokatzky, Margarida C. Gomes, Sydney L. Miles, Damián Lobato-Márquez, John Rohde, Jonathan N. Pruneda, Francis Impens, Serge Mostowy

## Abstract

Host cells contest invasion by intracellular bacterial pathogens with multiple strategies that recognise and / or damage the bacterial surface. To identify novel host defence factors targeted to intracellular bacteria, we developed a versatile proximity biotinylation approach coupled to quantitative mass spectrometry that maps the host-bacterial interface during infection. Using this method, we discovered that intracellular *Shigella* and *Salmonella* become targeted by UFM1-protein ligase 1 (UFL1), an E3 ligase that catalyses the covalent attachment of Ubiquitin-fold modifier 1 (UFM1) to target substrates in a process called UFMylation. We show that *Shigella* antagonises UFMylation in a dual manner: first, using its lipopolysaccharide (LPS) to shield from UFL1 recruitment; second, preventing UFM1 decoration by the bacterial effector IpaH9.8. Absence of UFMylation leads to an increase of bacterial burden in both human cells and zebrafish larvae, suggesting that UFMylation is a highly conserved antibacterial pathway. Contrary to canonical ubiquitylation, the protective role of UFMylation is independent of autophagy. Altogether, our proximity mapping of the host-bacterial interface identifies UFMylation as an ancient antibacterial pathway and holds great promise to reveal other cell-autonomous immunity mechanisms.

## Main

*Shigella flexneri* is a human-adapted enteroinvasive pathogen that causes over 200 million episodes of bacillary dysentery annually worldwide^1^. Its virulence relies on a Type 3 Secretion System (T3SS) that injects around 30 bacterial effectors into host cells to modulate key aspects of the infection process^2,3^. Once in the gastrointestinal tract, *S. flexneri* invades colonic epithelial cells and escapes from the phagosome to the cytosol within 10 minutes^4^. *S. flexneri* has evolved to thrive in the host cytosol, where it replicates and hijacks the actin cytoskeleton to form distinctive actin tails that enable intracellular motility and cell to cell spread^5,6^.

Cells protect their nutrient-rich cytosol from bacterial intruders with a plethora of highly conserved mechanisms collectively known as cell-autonomous immunity^7^. These include the prevention of actin tail motility by the entrapment of *S. flexneri* in septin cage-like structures^8,9^ and guanylate-binding protein (GBP) coats^10,11^. GBPs have also been shown to bind and disrupt the bacterial lipopolysaccharide (LPS)^12^, which exposes bacterial membranes to permeabilisation by Apolipoprotein L3^13^. In addition, bacterial LPS becomes ubiquitylated by the E3 ubiquitin ligase ring finger protein 213 (RNF213), which is required for the subsequent extension of the ubiquitin bacterial coat by other host E3 ligases (LRSAM1, Parkin, LUBAC and SMURF1)^14^. Although bacterial ubiquitylation by E3 ligases typically targets intracellular bacteria such as *Salmonella* to degradation by autophagy (xenophagy) via specific cargo receptors, *S. flexneri* avoids autophagosome recruitment by the effectors IcsB, VirA, and likely other mechanisms yet to be understood^15–20^.

In addition to canonical ubiquitylation, eukaryotic cells possess multiple ubiquitin-like (UBL) systems. These are post-translational modification machineries where small peptides with low sequence homology but high structural similarity to ubiquitin (including the β-grasp fold or ubiquitin fold) become conjugated to target substrates by the sequential action of an E1 activating, an E2 conjugating and an E3 ligating enzyme^21^, in a similar process to ubiquitylation. Although several UBL systems (FAT10ylation^22^, ISGylation^23^, NEDDylation^24,25^, SUMOylation^26,27^) are protective against various bacterial infections, none of them have been shown to have a direct antibacterial role at the bacterial surface analogous to ubiquitylation.

Here, we develop a versatile nanobody-based proximity biotinylation approach to map host proteins enriched in the vicinity of cytosolic *S. flexneri* during infection using quantitative mass spectrometry. Using this method, we discover that *S. flexneri* and *Salmonella enterica* subsp. Enterica srv. Typhimurium become targeted by UFMylation, a highly conserved UBL system^28^ that has recently emerged as a major regulator of cell homeostasis. This is particularly well studied at the endoplasmic reticulum (ER), where UFMylation is involved in stress tolerance through ribosome-associated quality control^29^ and selective ER autophagy (ER-phagy)^30^. In addition, UFMylation has been shown to be involved in DNA damage responses^31^, telomere length maintenance^32^, lipid droplet biogenesis^33^, autophagy regulation^34^, and immune signalling^35,36^. Here, we show that UFMylation also has a cell-autonomous immunity role against intracellular bacteria. We show that *S. flexneri* counteracts UFMylation via two distinct mechanisms: its LPS hampers the recruitment of the E3 ligase UFL1, and the secreted effector IpaH9.8 antagonises decoration of the UBL molecule UFM1 on the surface of intracellular bacteria. While UFMylated bacteria are not targeted to degradation by autophagy, lack of UFMylation leads to increased bacterial burden in human epithelial cells and zebrafish larvae. Together, we identify UFMylation as a highly conserved cell-autonomous immunity pathway against bacterial infection.

## Results

### Proximity biotinylation approach at the host-bacterial interface identifies UFL1 as targeted to intracellular bacteria

Recognition and restriction of intracellular pathogens by cell-autonomous immunity provides an immediate and localised defence against infection before the organism can mount an immune response. Importantly, many cell-autonomous immunity mechanisms that host cells deploy against bacterial intruders are delivered to the bacterial surface. We exploited this feature to develop a proximity biotinylation approach that specifically maps host proteins at the interface between the host and the bacterial surface during *S. flexneri* infection (i.e., the bacterial “proxisome”). For this, we engineered *S. flexneri* to display anti-GFP nanobodies on its surface, which enabled their functionalisation in vitro with GFP-fused recombinant proteins such as APEX2, an engineered ascorbate peroxidase widely used for proximity biotinylation campaigns^37^ (**Fig. 1a** and **Extended Data Fig. 1a, b**). In a second step, bacteria functionalised with GFP-APEX2 can be used in infection assays for the biotinylation of host proteins in close proximity to bacteria and subsequent identification by mass spectrometry. In vitro, we observed specific biotinylation of the *S. flexneri* surface when the anti-GFP nanobody was expressed and exogenous biotin-phenol and H_2_O_2_ were added (**Extended Data Fig. 1c-e**). We showed that GFP-APEX2 functionalised bacteria were able to infect HeLa cells and form actin tails and septin cages, which are considered hallmarks of the *S. flexneri* cytosolic lifestyle (**Extended Data Fig. 2a, b**). In HeLa, biotinylated proteins appeared as a halo around the bacteria during infection in immunofluorescence (**Fig. 1b**) and as a smear during western blot (**Fig. 1c**). Biotinylation of the *S. flexneri* microenvironment also occurred in vivo during infection of zebrafish larvae, highlighting the versatility of the method beyond tissue culture cells (**Fig. 1d** and **Supplementary Video 1**).

**Figure 1.**
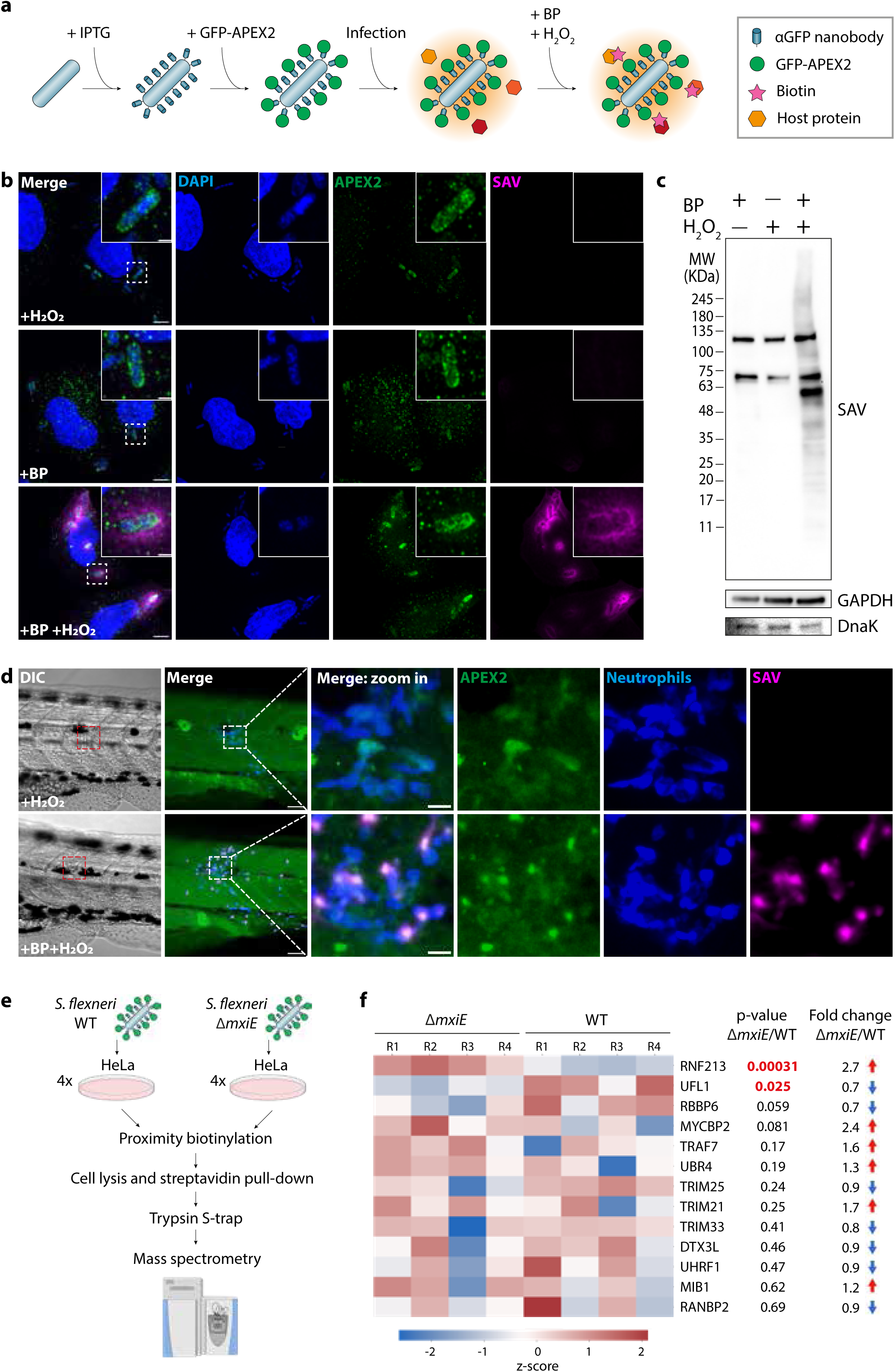
Novel proximity biotinylation approach to identify the *S. flexneri* proxisome during infection. **a,** Strategy for the in vitro functionalisation of *S. flexneri* for proximity mapping. First, an anti-GFP nanobody is displayed on the *S. flexneri* surface by induction with IPTG. After that, *S. flexneri* is coated with purified GFP-APEX2 in vitro. Then, functionalised bacteria are used for infection assays. Finally, the addition of biotin phenol (BP) and H_2_O_2_ enables the proximity biotinylation reaction during infection. **b,** Representative airyscan confocal images showing biotinylation in the vicinity of functionalised *S. flexneri* during infection in HeLa cells, in the presence or absence of BP and H_2_O_2_. SAV stands for streptavidin. Scale bar, 5 µm and 1 µm for the inset. **c,** Western blot shows a smear of biotinylated proteins in HeLa cells infected with functionalised *S. flexneri* upon addition of BP and H_2_O_2_. **d,** Proximity biotinylation occurs in zebrafish larvae at 45 minutes post infection of functionalised *S. flexneri* at the tail musculature. The transgenic zebrafish line *Tg(lyzC:DsRed2)^nz^*^50^ was used to visualise fluorescent neutrophils. Scale bar, 50 µm and 10 µm for the inset. **e,** Experimental workflow to identify the proxisome of *S. flexneri* WT and Δ*mxiE* mutant using mass spectrometry. Image created using BioRender. **f,** Heatmap of E3 ligases found in the proxisomes of *S. flexneri* during infection, inversely ordered by p-value.

Using this approach, we identified the proxisome of *S. flexneri* during infection in HeLa cells by label-free quantitative mass spectrometry. We compared the proxisomes generated by wildtype (WT) bacteria and bacteria lacking the transcriptional regulator MxiE, which controls expression of late T3SS effectors including important factors that antagonise cell-autonomous immunity (**Fig. 1e** and **Extended Data Fig. 2c, d**). To this end, HeLa cells were infected with GFP-APEX2 functionalised WT or Δ*mxiE S. flexneri* and biotinylated proteins were pulled-down for liquid chromatography-tandem mass spectrometry (LC-MS/MS analysis, **Fig. 1e**). We identified and quantified a total of 1.298 proteins of which 9 were *S. flexneri* proteins and 1.289 were human host proteins (**Supplementary Table 1** and **Extended Data Fig. 3**). GO term enrichment analysis of the host proteins identified multiple pathways related to host-pathogen interactions relevant for *S. flexneri* infection, including actin filament-based transport, outer-mitochondrial membrane organisation or stress granule assembly (**Extended Data Fig. 3c**). In addition, proteomic screening revealed that multiple members of the ubiquitylation machinery are recruited to the bacterial surface during infection, including 13 E3 ligases (**Fig. 1f**, and **Extended Data Fig. 3d**). One of the most enriched hits in the proxisome of the *S. flexneri* Δ*mxiE* was the E3 ligase ring finger protein 213 (RNF213), which was discovered to ubiquitylate the LPS of cytosolic *S.* Typhimurium^14^ and *S. flexneri*^18–20^. It has been recently shown that the MxiE*-*dependent effector IpaH1.4 prevents RNF213 recruitment to *S. flexneri*, explaining why RNF213 is enriched to Δ*mxiE* compared to WT bacteria in our proximity data. Considering the pivotal role of ubiquitin E3 ligases in host defence, we decided to focus on the E3 UFM1-protein Ligase 1 (UFL1, **Fig. 1f**), an enzyme that catalyses the covalent attachment of Ubiquitin-like modifier 1 (UFM1) to target proteins in a process called UFMylation similar to ubiquitylation. UFL1 had the second lowest p-value among the E3 ligases identified (**Fig. 1f**). Together, we show that our proximity biotinylation approach is effective to identify novel host factors in close proximity to *S. flexneri* during infection, including UFL1 and other E3 ligases.

### *S. flexneri* LPS shields intracellular bacteria from recognition by the UFMylation ligase UFL1

Microscopy experiments validated the localisation of UFL1 and UFM1 to the surface of *S. flexneri* during infection in HeLa cells, using both HA-tagged exogenous expression and antibodies against the endogenous proteins (**Fig. 2a-c** and **Extended Data Fig. 4**). Considering that LPS is the bacterial substrate for ubiquitylation by the E3 ligase RNF213^14^, we tested the impact of UFL1 and UFM1 recruitment to a *S. flexneri* mutant Δ*rfaC*, which lacks the O-antigen and the LPS outer core^38^. We observed a strong increase in UFL1 recruitment in *S. flexneri* Δ*rfaC* (2.1 ± 0.2-fold, **Fig. 2a, b** and **Extended Data Fig. 5d**), suggesting that LPS protects intracellular bacteria from UFL1 binding. However, this increase in UFL1 binding did not correlate with an increase in UFM1 decoration of *S. flexneri* (WT 12.2% ± 5.0%, Δ*rfaC* 13.1% ± 8.5% bacteria, **Fig. 2c** and **Extended Data Fig. 5e**), suggesting that additional bacterial factors may prevent bacterial UFMylation. In both *S. flexneri* WT and Δ*rfaC* we observed similar and low values of ubiquitylation (WT: 3.9% ± 2.9 and 6.6 ± 3.6 bacteria, respectively), and there was little overlap between UFM1 and ubiquitin positive bacteria (**Figure 2c**), consistent with IpaH1.4 promoting RNF213 degradation^18–20^.

**Figure 2.**
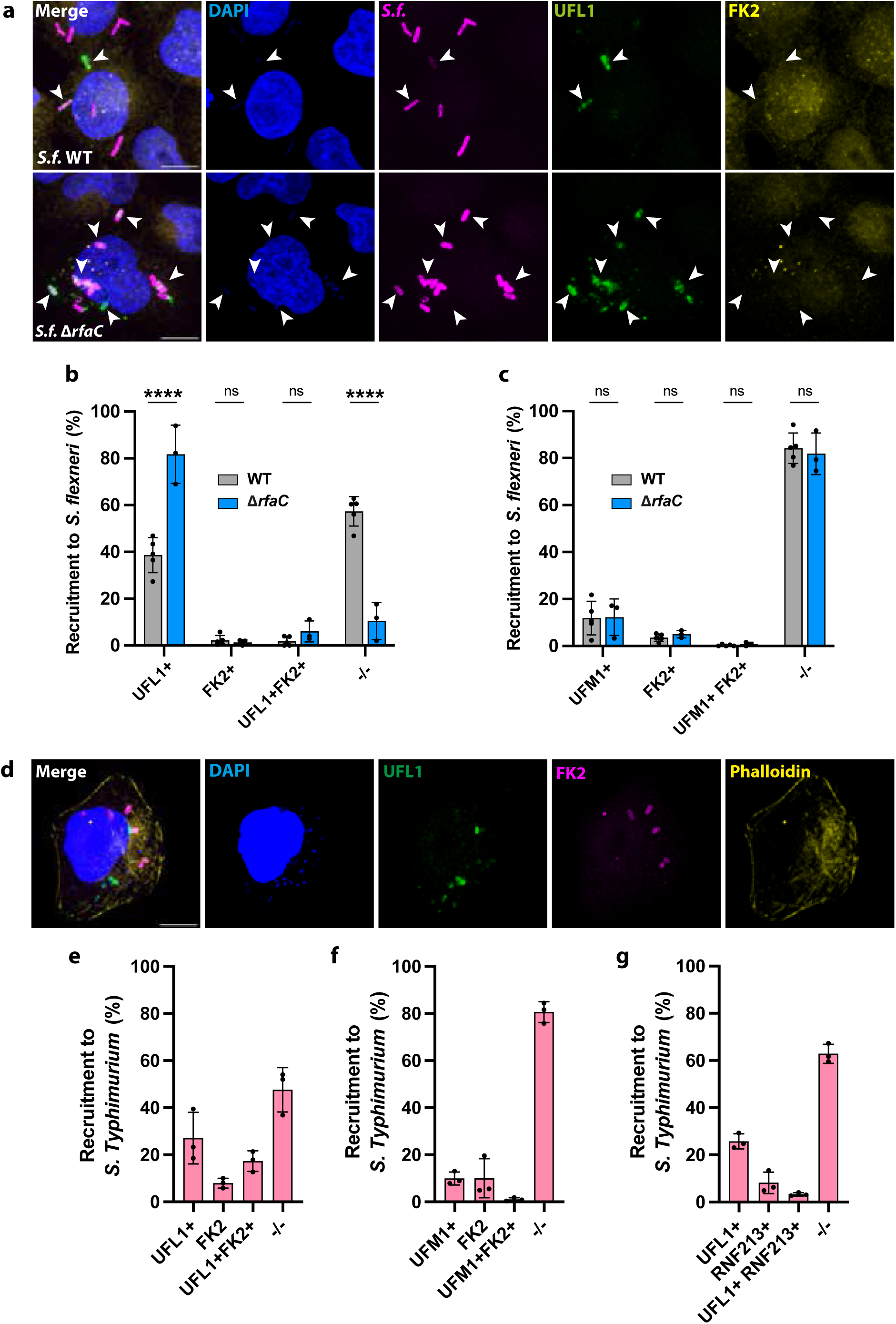
Several members of the UFMylation pathway are recruited to *S. flexneri* and *S.* Typhimurium during infection. **a,** Representative confocal images of HeLa cells infected with *S. flexneri* WT and Δ*rfaC*. Arrows show UFL1 recruitment to intracellular bacteria. Scale bar, 10 µm. **b,** Percentage of *S. flexneri* WT (n=1,311) and Δ*rfaC* (n=630) colocalising with UFL1 and ubiquitin (FK2), from at least 3 biological replicates. Two-way ANOVA and Tukey’s multiple comparison test. **c,** Percentage of *S. flexneri* WT (n=882) and Δ*rfaC* (n=643) colocalising with UFM1 and ubiquitin (FK2), from at least 3 biological replicates. Two-way ANOVA and Tukey’s multiple comparison test. **d,** Representative confocal image of HeLa cell infected with *S.* Typhimurium and stained for UFL1 and ubiquitin (FK2). Scale bar, 10 µm. **e,** Percentage of *S.* Typhimurium (n=740) colocalising with UFL1 and ubiquitin (FK2) **f,** Percentage of *S.* Typhimurium (n=823) colocalising with UFM1 and ubiquitin (FK2). **g,** Percentage of *S.* Typhimurium (n=735) colocalising with UFL1 and GFP-RNF213. The results are represented as mean ± SD.

We hypothesised that the UFMylation machinery may recognise other intracellular bacteria beyond *S. flexneri*. We tested *S.* Typhimurium, a bacterium that typically resides within vacuoles but occasionally escapes into the cytosol, where it hyperreplicates^39^. We observed that UFL1 and UFM1 were also recruited to the surface of *S.* Typhimurium at 3 hours post infection (**Fig. 2d-g**), indicating that bacterial UFMylation is not a phenotype exclusive to *S. flexneri*. Most bacteria recruiting UFL1 or UFM1 did so in the absence of GFP-RNF213 recruitment or ubiquitylation (**Fig. 2d-g** and **Extended Data Fig. 5f**). Despite some overlap between UFL1 and FK2 decoration (17.3% ± 4.4% of the bacteria), robust recruitment of UFL1 and FK2 rarely coincided (**Fig. 2d**). Together, these results suggest that UFMylation and ubiquitylation are functionally separate machineries.

### UFMylation of intracellular *S. flexneri* is antagonised by the bacterial effector IpaH9.8

The limited UFM1 decoration of *S. flexneri* during infection led us to hypothesise that additional bacterial effectors may intersect with the UFMylation cascade beyond UFL1 recruitment. Among T3SS effectors, members of the Invasion Plasmid Antigen H (IpaH) family of proteins have been shown to have important roles in counteracting cell-autonomous immunity mechanisms^40–42^. Importantly, a Yeast-Two-Hybrid screening for the *S. flexneri* secreted effector IpaH9.8 identified UFM1 as potential interaction partner (**Supplementary Table 2**), suggesting that IpaH9.8 may prevent *S. flexneri* UFMylation. Immunoprecipitation assays using recombinantly expressed IpaH9.8 enhanced the recovery of UFM1 species (**Extended Data Fig. 6**). Binding studies using purified proteins confirmed a weak but direct interaction between GST-tagged IpaH9.8 and UFM1 (**Fig. 3a**). Meanwhile, as a control, GST-tagged ubiquitin showed no interaction with UFM1. To understand the interaction of *S. flexneri* with UFM1, we isolated bacteria from infected cells and probed for the presence of UFM1 conjugates by western blot (**Fig. 3b**). We detected several UFM1 positive bands that were absent in bacteria from broth, indicating UFM1 conjugates at the bacterial surface. Interestingly, we observed a similar pattern in a *S. flexneri* double mutant Δ*rfaC*Δ*ipaH9.8* (**Fig. 3b** and **Extended Data Fig. 5a, c**), suggesting that LPS is not the bacterial substrate for UFMylation and that IpaH9.8 does not prevent the conjugation of specific substrates. To test a role for IpaH9.8 in counteracting *S. flexneri* UFMylation, we scored UFL1 and UFM1 recruitment to *S. flexneri* WT, Δ*ipaH9.8* and the double mutant Δ*rfaC*Δ*ipaH9.8*. While deletion of *ipaH9.8* did not affect UFL1 recruitment to *S. flexneri*, we observed a higher decoration of UFM1 in *S. flexneri* Δ*ipaH9.8* compared to WT at 3 hours post infection (2.6 ± 0.5-fold), which further increased (4.4 ± 0.6-fold) in the case of the double mutant Δ*rfaC*Δ*ipaH9.8* (**Fig. 3c-e**, and **Extended Data Fig. 5a-e)**. At later timepoints (5 hours post infection) the levels of UFM1 decoration were higher and non-significantly different for *S. flexneri* WT, Δ*ipaH9.8* and the double mutant Δ*rfaC*Δ*ipaH9.8* (37.8% ± 30.6%, 54.6 ± 26.7%, 60.6% ± 12.1%, respectively, **Extended Data Fig. 5a-e**). Consistent with previous reports^10^, deletion of *ipaH9.8* in the absence of IFNγ did not impact *S. flexneri* ubiquitylation (**Fig. 3d, e**), further suggesting that UFMylation operates independently of the ubiquitin axis. Together, these results indicate that the *S. flexneri* T3SS effector IpaH9.8 interacts with UFM1 to delay UFMylation of the bacterial surface during infection.

**Figure 3.**
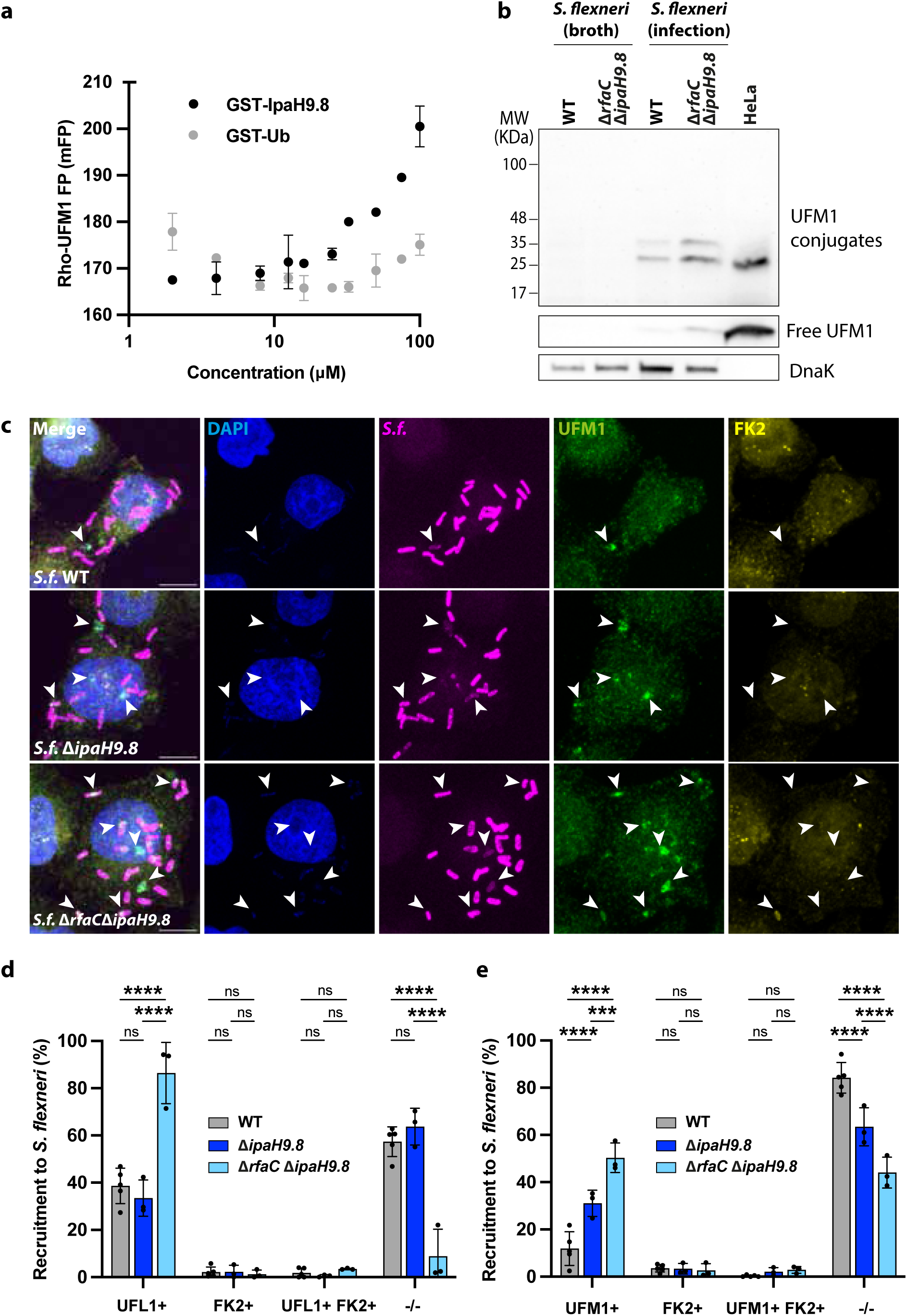
The *S. flexneri* secreted effector IpaH9.8 prevents UFM1 conjugation of intracellular bacteria. **a,** Protein-protein interaction assay monitoring fluorescence polarisation (FP) of rhodamine-labelled UFM1 in response to increasing concentrations of recombinant GST-IpaH9.8. To control for possible interactions with the GST tag of GST-IpaH9.8, GST-ubiquitin is titrated instead. **b,** Western blot showing UFM1 conjugates in *S. flexneri* WT and Δ*rfaC*Δ*ipaH9.8* purified from infection that are absent in bacteria from broth. UFM1 species present in uninfected HeLa cells are shown as a control. Different exposure times were used to detect free UFM1 (low) and UFM1 conjugates (high) due to relative differences in abundance, using anti-UFM1 antibodies (Abcam). **c,** Representative confocal images of HeLa cells infected with *S. flexneri* WT, Δ*ipaH9.8* and Δ*rfaC*Δ*ipaH9.8*. Arrows show UFL1 recruitment to intracellular bacteria. Scale bar, 10 µm. **d,** Percentage of *S. flexneri* WT (n=1,311), Δ*ipaH9.8* (n=463) and Δ*rfaC*Δ*ipaH9.8* (n=553) colocalising with UFL1 and ubiquitin (FK2). **e,** Percentage of *S. flexneri* WT (n=882), Δ*ipaH9.8* (n=476) and Δ*rfaC*Δ*ipaH9.8* (n=526) colocalising with UFM1 and ubiquitin (FK2). Two-way ANOVA and Tukey’s multiple comparison test. UFL1 and UFM1 quantifications for WT *S. flexneri* are the same as presented in Fig. 2b, c. The results are represented as mean ± SD.

### UFMylation has a conserved antibacterial role independent of autophagy

Considering the ubiquitous expression of the UFMylation pathway in human cells (Human Cell Atlas) and its high conservation among eukaryotes^28^, we decided to test the localisation of UFL1 and UFM1 to *S. flexneri* in THP-1 macrophages and zebrafish larvae. In both cases, UFL1 and UFM1 localised to *S. flexneri* (**Extended Data Fig. 7**), suggesting a broad and conserved role during infection.

We next tested whether UFMylation has an antibacterial role, similar to ubiquitylation. We observed that intracellular *S. flexneri* with a strong recruitment of UFL1 or UFM1 displayed a fluorescence loss of their mCherry constitutive reporter, suggesting UFMylated bacteria were subjected to degradation (**Fig. 4a**). Similar loss of fluorescence was also observed in THP-1 macrophages and zebrafish larvae (**Extended Data Fig. 7**). Considering ubiquitylation classically targets intracellular bacteria to degradation by the autophagic machinery, we assessed the colocalisation of UFMylation and the main autophagic marker GFP-LC3 during infection. In the case of *S. flexneri*, we observed no overlap between UFL1 recruitment and GFP-LC3 engulfment of intracellular bacteria (**Fig. 4b, c**), consistent with ubiquitylation results. *S. flexneri* mutants with increased UFL1 and/or UFM1 colocalisation (Δ*ipaH9.8*, Δ*rfaC*, Δ*rfaC*Δ*ipaH9.8*) did not increase the number of bacteria in autophagosomes (**Fig. 4c**). In the case of *S.* Typhimurium, most of the bacteria entrapped in an autophagosome also did not recruit UFL1 (**Fig. 4d**). Together, these results suggest that contrary to ubiquitylation, UFMylation of intracellular bacteria leads to bacterial degradation in a process independent of autophagy.

**Figure 4.**
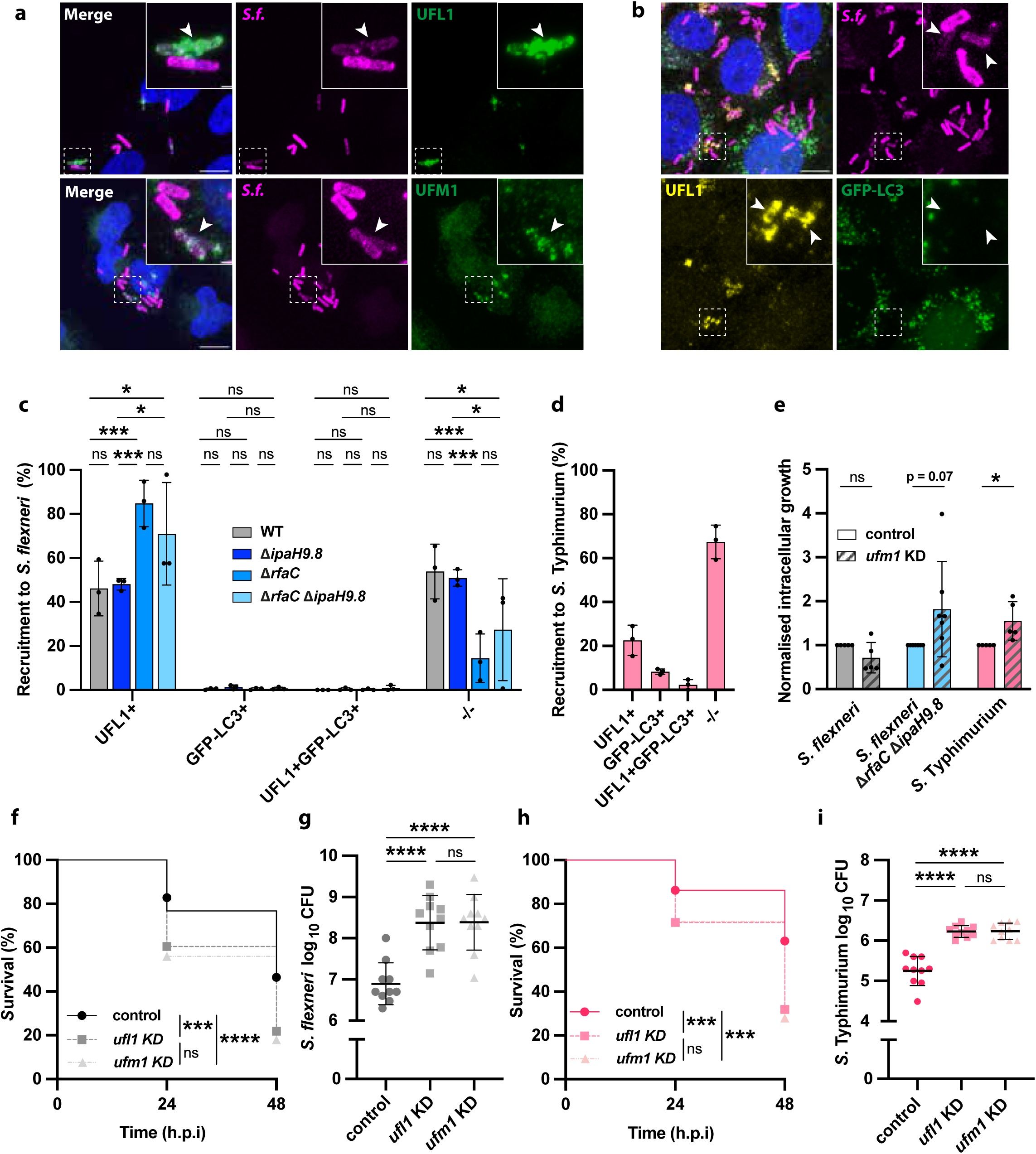
UFMylation has a conserved antibacterial role independent of autophagy. **a,** Representative confocal microscopy images of HeLa cells infected with *S. flexneri* mCherry and immunostained against UFL1 and UFM1. Arrows point at intracellular bacteria colocalising with UFL1 and UFM1 with a loss of mCherry fluorescence. Scale bar, 10 μm and 1 μm for the inset. **b,** Representative microscopy image of HeLa cells expressing GFP-LC3 infected with *S. flexneri* mCherry and immunostained against UFL1. Arrows indicate intracellular *S. flexneri* that recruit UFL1 and lose their mCherry fluorescence but are not engulfed in an autophagosome. Scale bar, 10 μm and 1 μm for the inset. **c,** Quantification of UFL1 and GFP-LC3 recruitment to *S. flexneri* WT (n=655), Δ*ipaH9.8* (n=733), Δ*rfaC* (n=422), and double Δ*rfaC*Δ*ipaH9.8* (n=627) mutants. Two-way ANOVA and Tukey’s multiple comparison test. **d,** Quantification of UFL1 and GFP-LC3 recruitment to *S.* Typhimurium (n=582). **e,** Intracellular growth of *S. flexneri* WT, Δ*rfaC*Δ*ipaH9.8* and *S.* Typhimurium in HeLa cells treated with *ufm1* siRNA normalised to control siRNA after 5 hours post infection. **f,** Zebrafish larvae survival after *S. flexneri* infection at the caudal vein for wildtype larvae (46.40%) and *ufl1* (21.85%) and *ufm1* (18.00%) KD crispants. Data was obtained from 76 wildtype, 81 *ufl1* and 70 *ufm1* KD crispant zebrafish larvae from two biological replicates. **g,** *S. flexneri* recovered from infected zebrafish larvae at 48 hours post infection (h.p.i) for wildtype larvae, *ufl1* and *ufm1* KD crispants. **h,** Zebrafish larvae survival after *S.* Typhimurium infection at the caudal vein for wildtype larvae (63.15%), *ufl1* (31.81%) and *ufm1* (27.91%) KD crispants. Data was obtained from 54 wildtype, 61 *ufl1* and 55 *ufm1* KD crispant zebrafish larvae from two biological replicates. **i,** *S.* Typhimurium recovered from infected zebrafish larvae at 48 hours post infection, for wildtype larvae, *ufl1* and *ufm1* KD crispants. The results are represented as mean ± SD.

To further investigate the antibacterial role of the UFMylation pathway, we assessed its impact on *S. flexneri* and *S.* Typhimurium burden in cells (**Fig. 4e** and **Extended Data Fig. 8)**. Depletion of *ufm1* using siRNA did not lead to an increase in intracellular bacterial replication in the case of *S. flexneri*, consistent with its capacity to antagonise the UFMylation pathway. *S. flexneri* Δ*rfaC*Δ*ipaH9.8* exhibited more pronounced growth upon *ufm1* depletion, but it was non-significant (p-value =0.070), suggesting that additional effectors may counteract this pathway. In the case of *S.* Typhimurium, a vacuolar pathogen with a limited arsenal for cytosolic survival, we observed a significant increase in intracellular replication upon *ufm1* knockdown (1.5 ± 0.4-fold, **Fig. 4e** and **Extended Data Fig. 8**). Together with the loss of fluorescence of UFMylation targeted cells, these results indicate that UFMylation plays a host defence role against cytosolic bacteria.

Finally, we decided to interrogate the role of the UFMylation pathway in vivo using the zebrafish infection model. We observed a strong decrease in zebrafish larvae survival during infection upon depletion of *ufm1* and *ufl1*, both in the case of *S. flexneri* and *S.* Typhimurium (**Fig. 4f, h** and **Extended Data Fig. 9**). This increased susceptibility correlated with increased bacterial load of more than an order of magnitude in both cases (**Fig. 4g, i**). Collectively, these results provide evidence for an important and conserved defence role against bacterial infection and highlight the UFMylation pathway as a conserved / ancient cell-autonomous immunity mechanism (**Extended Data Fig. 10**).

## Discussion

Proximity mapping techniques have emerged as key discovery tools in a wide variety of research fields. For infection biology, recent applications include screenings for novel host interacting partners of *Legionella*^43^ and *S.* Typhimurium^44^ secreted effectors, as well as the surface proteome (including host bound proteins) of *Vibrio cholerae* recovered from rabbit diarrheal fluid^45^. Here, we pioneered a proximity biotinylation approach to specifically map the host-bacterial interface during infection. This strategy is particularly relevant to understand the proteomic microenvironment of cytosol-dwelling bacteria, since the isolation and proteomic characterisation of bacterial containing vesicles (BCVs) has been performed successfully for multiple vacuolar pathogens^46–50^. In addition, our proximity biotinylation approach is highly versatile, as it is functional in tissue culture cells and in animal models, and it can be easily adaptable to any other genetically tractable Gram positive or negative bacteria.

Our proximity biotinylation screening identified multiple E3 ligases in close proximity to *S. flexneri* during infection. Among these, RNF213 was found enriched in bacteria lacking MxiE, an important transcriptional regulator for the expression of late T3SS effectors. Consistent with these results and serving as internal positive control for our screening, multiple studies have recently reported that *S. flexneri* deploys the bacterial effector IpaH1.4 (under MxiE transcriptional control) to target RNF213 to degradation^18–20^. The proxisome of *S. flexneri* also revealed UFL1, the only identified E3 ligase of the UFMylation pathway, the latest UBL system discovered.

Our work has shown that both UFL1 and its substrate peptide UFM1 are recruited to the surface of *S. flexneri* and *S.* Typhimurium during infection, which showed little to no overlap with bacterial ubiquitylation. The *S. flexneri* surface became UFMylated during infection, as bacteria recovered from infected cells presented UFM1-conjugates absent in bacteria from broth. In contrast to RNF213, which ubiquitylates LPS^14^, the nature of the UFMylated substrates on *S. flexneri* is likely proteinaceous, as they appeared as discrete, well-defined bands on western blot. This result indicates that bacterial ubiquitylation by RNF213 and UFMylation by UFL1 are separate pathways that protect the host cytosol in a multilayered manner.

*S. flexneri* has evolved strategies to avoid UFMylation, which underscores its relevance as an antibacterial pathway. First, the S*. flexneri* mutant Δ*rfaC* with shortened LPS showed an increased recruitment of UFL1, indicating that the LPS can shield intracellular bacteria from host recognition. It is worth noting that *S. flexneri* Δ*rfaC* is also more susceptible to septin cage entrapment^38^. Collectively, these results suggest that the protective role of the LPS may contribute not only to avoid extracellular recognition^51,52^, but also intracellular detection. Likely reflecting the host-pathogen arms race, host cells also evolved to recognise intracellular LPS by caspase-4/-11^53^ and the TRAF-interacting forkhead-associated protein A (TIFA)^54,55^ for inflammasome and NF-κB activation, respectively. In this context, it is not surprising that bacterial pathogens, including *S. flexneri*, tightly modulate their LPS production during infection^56,57^.

A second strategy deployed by *S. flexneri* to counteract UFMylation is the secreted effector IpaH9.8, which did not impact UFL1 recruitment to the bacteria but their decoration with UFM1. This adds to the list of known functions of IpaH9.8, which has been shown to dampen the host inflammatory response^58,59^ and interfere specifically with established cell-autonomous immune programs by targeting GBPs to proteasomal degradation^10,11^. It is remarkable how IpaH9.8 has evolved for such level of pleiotropy, and future structural and biochemical studies will delve into the specificity and molecular mechanism underlying the IpaH9.8 – UFM1 interaction.

Our work has revealed UFMylation as a novel cell-autonomous immune pathway against cytosolic bacteria. This pathway is active during infection of human cells (epithelial and THP-1 macrophages) and also in zebrafish larvae. Together with the high degree of conservation of the UFMylation cascade among eukaryotes^28^, this strongly suggests a broader antibacterial role across evolution. Remarkably, UFMylation has very recently been shown to participate in the regulation of interferon and NF-κB mediated responses during viral and bacterial infections^35,60–63^, which may support the different susceptibility to *S. flexneri* and *S.* Typhimurium observed in cellulo and in vivo. Taken together, the UFMylation machinery should now be recognised as a major contributor to host defence.

## Acknowledgements

We thank members of the Mostowy lab for helpful discussion and the LSHTM Biological Services Facility for the work and care of zebrafish stocks. We thank the VIB Proteomics Core for help with the mass spectrometry analyses. We thank Luis Ángel Fernández from Centro Nacional de Biotecnología (CNB-CSIC) for sharing the plasmids for the display of nanobodies on the bacterial surface and Prof. Terje Johansen from the Artic University of Norway for sharing the GFP-LC3 expressing HeLa cell line. We thank Dr. Craig McCormick and Carolyn Robinson from Dalhousie University for construction and gift of plasmid pCR3.1-His-UFM1. A.T.L.J. was funded by the Swiss National Science Foundation Early Postdoc.Mobility Fellowship (P2GEP3_188277) and the European Union’s Horizon 2020 research and innovation program under the Marie Skłodowska - Curie grant agreement no. H2020-MSCA-IF-2020-895330. F.T. acknowledges support by the Research Foundation-Flanders (FWO) through postdoctoral fellowship 12AN524N. Research in the J.N. Pruneda laboratory was supported by the National Institute of General Medical Sciences (R35GM142486). Research in the F. Impens laboratory was supported by a Ghent University Concerted Research Action grant BOF21/GOA/033 and a Starting Grant BOF/STA/202209/011, as well as by the European Research Council (ERC Consolidator Grant #101089193). Research in the S. Mostowy laboratory was supported by a Wellcome Trust Senior Research Fellowship (206444/Z/17/Z), European Research Council Consolidator Grant (772853 - ENTRAPMENT) and Wellcome Discovery Award (226644/Z/22/Z).

## Methods

### Reagents

The following primary antibodies were used: mouse anti-Ubiquitinylated proteins, clone FK2 (Sigma, #04-263), rabbit anti-UFL1 (Proteintech, #26087-1-AP), rabbit anti-SEPT7 (IBL, #18991), mouse anti-HA-Tag (6E2) (Cell Signaling, #2367), rabbit anti-Myc-Tag (71D10) (Cell Signaling, #2278), mouse anti-GFP (Abcam, #ab1218) and rabbit anti-IpaH^64^. Rabbit anti-UFM1 (Proteintech, #15883-1-AP) was used for immunofluorescence, and rabbit anti-UFM1 (Proteintech, #15883-1-AP; BostonBiochem, #AF8237; Abcam #109305) were used for western blot as indicated in the figure legends. The following secondary antibodies were used: Alexa Fluor-488, −555 and −647 conjugated goat anti-mouse antibodies (Invitrogen #A-11001, #A-21424 and #A-21236), and Alexa Fluor-488, −555 and −647 conjugated goat anti-rabbit antibodies (Invitrogen, #A-11008, #A-21428, # A-21244). The following reagents were used: HRP-conjugated streptavidin (Invitrogen, #S911), Alexa Fluor-488 conjugated streptavidin (Invitrogen, #S32354), Alexa Fluor-647 conjugated phalloidin (Invitrogen, #A22287) Congo red (Sigma-Aldrich, #C6767) and IFNγ (R&D Systems, #285-IF).

### Bacterial strains and culture conditions

The bacterial strains and plasmids described in this study are listed in **Supplementary Table 3**. *Shigella flexneri* 5a str. M90T was grown in trypticase soy broth (TCS) agar containing 0.01% of Congo red to select for red colonies, indicative of a functional T3SS. Conical polypropylene tubes containing 5 ml of TCS were inoculated with individual red colonies of *S. flexneri* and were grown ∼16 h at 37°C with shaking at 200 rpm. The following day, bacterial cultures were diluted in fresh prewarmed TCS (1:50 or 1:100 v/v), and cultured until an optical density (OD, measured at 600 nm) of 0.6. For *Salmonella enterica* subs. *enterica* srv. Typhimurium str. SL1344, single bacterial colonies were grown in conical polypropylene tubes containing 5 mL of LB for ∼16 h at 37°C in static conditions. The following day, bacteria were subcultured in fresh prewarmed LB (1:50 or 1:100 v/v) and grown for 3h.

### Mammalian cell culture

HeLa (ATCC #CCL-2) and HEK293T (ATCC #CRL-3216) were cultured in Dulbecco’s modified Eagle Medium (DMEM, GIBCO) in the presence of 10% heat-inactivated fetal bovine serum (hi-FBS, Gibco, #10500064) under standard conditions (37°C and 5% CO_2_). GFP–LC3B-producing HeLa cells^65^ were grown as above.

THP-1 cells (male monocytes, ATCC #TIB-202) were grown RPMI-1640 medium supplemented with 10% hi-FBS at 37°C and 5% CO_2._ THP-1 cells were differentiated via phorbol 12-myristate 13-acetate (PMA, Sigma-Aldrich, #P8139) treatment^66^ into macrophages using 100 nM PMA in RPMI-1640 with 10% hi-FBS for 48 h followed by 24 h recovery phase in RPMI with 10% hi-FBS.

### Cloning

Plasmids used or generated in this study are listed in **Supplementary Table 4**. Primers used in this study were designed using Benchling (https://benchling.com) or NEBuilder Assembly Tool (https://nebuilder.neb.com/#!/) and are listed in **Supplementary Table 5**.

For GFP-APEX2 purification, the pTRC-GFP-APEX2 plasmid was generated by Gibson assembly using pTRC-APEX2^37^ and pLVX-msGFP–SEPT6^38^ as template.

For Yeast-Two-Hybrid experiments, DNA fragment encoding IpaH9.8 C337A was amplified by PCR from plasmid pJR006^40^ and cloned as an EcoRI-BamHI fragment into the vector pGKBT7 to create the Yeast-Two-Hybrid bait vector pJB179-10.

For immunoprecipitation experiments, DNA fragments encoding IpaH9.8 and UFM1 were amplified by PCR and cloned as EcoRI-BamHI fragments into the vectors pRK5-Myc and pCR3.1 respectively to create pRK5Myc-IpaH9.8 and pCR3.1His-UFM1.

*S. flexneri* mutants were engineered using λ-Red-mediated recombination^67^. In brief, kanamycin resistance-encoding DNA cassettes were amplified using pKD4 plasmid as template and primers containing 50 bp nucleotides homologous to the site of insertion. Resulting DNA fragments were electroporated in *S. flexneri* electrocompetent cells producing λ-Red recombinase and plated in TSA plates supplemented with 0.01% of Congo red and 50 μg/ml of kanamycin. All strains were verified by PCR as shown in **Extended Data Fig. 7**. *S.* Typhimurium was transformed with the pmCherry plasmid, containing an *mCherry* gene controlled under a *P_BAD_* arabinose-inducible promoter. *mCherry* was PCR-amplified from a pRK2-mCherry plasmid^68^ and cloned between ZraI and SpeI restriction sites in a pFUS-P_BAD_ plasmid^69^.

### siRNA treatment

HeLa cells were transfected using siRNA targeting UFM1 (Thermo Fisher scientific, #148645) or using negative control siRNA (Thermo Fisher scientific, #4390843). HeLa cells were seeded in a 6-well plate and 200 nM of siRNA was transfected per well using Oligofectamine (Invitrogen, #12252011) as described by the manufacturer. Cells were incubated for 48 hours to ensure sufficient knockdown of targeted proteins.

### DNA transfections

Hela and HEK273T cells were transfected using jetPEI transfection reagent (Polyplus, #101000053) according to the manufacturer’s instructions. Cells were incubated for 24 hours before the experiment.

### Protein purification

For the in vitro binding of APEX2 to anti-GFP nanobodies displayed on the surface of *S. flexneri*, a monomeric superfolder version of GFP was added between the His_6_ tag and the N-terminus of APEX2 in the pTRC-APEX2 plasmid. GFP-APEX2 was purified as described previously^37,70^. Briefly, *Escherichia coli* BL21-DE3 cells transformed with the pTRC-GFP-APEX2 plasmid were grown to an OD (600 nm) of 0.5 in LB supplemented with 10 μg/mL ampicillin at 37°C with shaking at 200 rpm. Protein expression was induced with 1 mM IPTG (Sigma-Aldrich, #I5502) and supplemented with 1 mM 5-aminolevulinic acid hydrochloride (Sigma-Aldrich, #A7793) overnight at room temperature to promote heme incorporation. Harvested cells were lysed using B-PER in the presence of 1 mM PMSF (Thermo Fisher scientific, #36978) and cOmplete™ Protease Inhibitor Cocktail (Roche, #4693116001). The clarified lysates were incubated with Ni-NTA agarose chromatography matrix (Qiagen, #30210) for 30 minutes and loaded on an Econo-Pac column (Bio-Rad, #7321010) using gravity flow at 4 °C. The matrix was washed with binding buffer (50 mM Tris-HCl, 300 mM NaCl, pH 7.8) and washing buffer (binding buffer with 30 mM imidazole). GFP-APEX2 was eluted in binding buffer containing 200 mM imidazole. Purified protein was dialysed in PBS at 4°C and concentrated using a 30 KDa Amicon Ultra centrifugal filter unit (Millipore, #Z648035).

Expression plasmids for GST-IpaH9.8 and GST-ubiquitin, both in the pGEX6P-1 vector, were kind gifts from Dr. Neal Alto and Dr. David Komander, respectively. Protein expression was performed as described above with the exception of using Rosetta *E. coli* induced with 0.2 mM IPTG overnight at 18 °C. Cells were resuspended in 25 mM Tris-HCl, 200 mM NaCl, 2 mM *β*-mercaptoethanol, pH 8.0 and lysed by sonication in the presence of lysozyme, DNase, and protease inhibitor cocktail. The clarified lysates were loaded onto GST agarose (Pierce) in an Econo-Pac column for 60 minutes at 4 °C. The matrix was washed with the lysis buffer and eluted in lysis buffer containing 50 mM glutathione. Purified protein was dialysed in 25 mM Tris-HCl, 200 mM NaCl, 2 mM *β*-mercaptoethanol, pH 8.0 at 4°C and concentrated using a 30 KDa Amicon Ultra centrifugal filter unit (Millipore, #Z648035).

### In vitro coating of bacteria with GFP-APEX2

*S. flexneri* WT or Δ*mxiE* mutant carrying pNVgfp plasmid for nanobody display on the bacterial surface^71^ and pAC-*afaI* plasmid to promote hyper invasion of HeLa cells^72^ were grown overnight in the presence of the appropriate antibiotics. The following day, bacteria cultures were diluted 1:50 and grown until OD (600 nm) 0.6 in the presence of the appropriate antibiotics and 1 mM of IPTG. 1.5 mL of bacterial culture was washed twice in PBS and incubated in 75 μL of PBS containing 300 nM of purified GFP-APEX2 at room temperature on rotation for 1 hour. Bacteria were washed twice with PBS to remove unbound GFP-APEX2 and used for subsequent infection assays.

### Proximity biotinylation

Proximity biotinylation was adapted from previously published protocols^73^. 6 μM hemin-Cl (Sigma-Aldrich, #H9039) was added to all samples for the duration of the experiment to increase functionality of purified GFP-APEX2. For the biotinylation reaction, 500 μM of biotin-phenol (Sigma-Aldrich, #SML2135) was added to the sample and incubated for 30 minutes. 1 mM of H_2_O_2_ (Sigma-Aldrich, #H1009) was added and samples were incubated at 37°C for 1 minute (in vitro assay, in cellulo assay), or for 5 minutes (for zebrafish larvae). The biotinylation reaction was stopped by washing the cells in ice chilled quencher buffer containing 10 mM sodium ascorbate (Sigma-Aldrich, #A92902), 5 mM trolox (Sigma-Aldrich, #238813) and 10 mM sodium azide (Thermo Fisher Scientific #190380050) three times. Cells were collected using a cell scraper for western blot analysis or purification of biotinylated proteins. For immunofluorescence, samples were first fixed with 4% paraformaldehyde (in PBS) at room temperature and proximity biotinylation was performed on fixed cells or zebrafish larvae to prevent diffusion of biotinylated proteins.

### Purification of biotinylated proteins

Purification of biotinylated proteins was performed as described before^73^. After proximity biotinylation, collected cells were centrifuged for 10 minutes at 3,000 xg at 4°C and lysed in RIPA buffer (750 μL for 10^7^ cells) supplemented with cOmplete™ Protease Inhibitor Cocktail, 1 mM PMSF and quenchers (10 mM sodium azide, 10 mM sodium ascorbate and 5 mM trolox), and incubated on ice for 2 minutes. Lysates were clarified by centrifugation at 15,000 xg for 10 minutes at 4 °C. Pierce™ Streptavidin magnetic beads (Thermo Fisher Scientific, #88816) were added to whole cell lysates (60 μL for 10^7^ cells) and incubated on rotation 1 hour at room temperature. A magnetic rack was used to pellet the beads and collect the supernatant (flow-through). Beads were washed to remove nonspecific binders in ice cold conditions twice with RIPA buffer, once with 1 M KCl, once with 0.1 M Na_2_CO_3_, once with 2 M urea in 10 mM Tris-HCl, pH 8.0, and twice with RIPA buffer again. Beads were snap frozen and subsequently analysed by mass spectrometry or alternatively processed for western blot analysis. For western blot, biotinylated proteins were eluted from the beads by boiling 50 μL of beads in 50 μL Laemmli buffer (3x) supplemented with 2 mM biotin and 20 mM DTT for 10 minutes and finally recovered from the beads using a magnetic rack.

### Sample preparation for LC-MS/MS analysis

4 x 10^7^ HeLa cells were treated with 1 ng/uL IFNγ for 16 hours and were infected with GFP-APEX2 coated S*. flexneri* WT or Δ*mxiE* at MOI 80 in quadruplicates. Proximity biotinylation was performed at 1 hour 30 minutes post infection and biotinylated proteins were purified. Streptavidin beads were resuspended with 23 µL of a buffer containing 5% SDS (Merck, #L4509) with 100 mM triethylammonium bicarbonate (TEAB) (Merck, #T7408) at pH 8.0. Proteins were reduced and alkylated with 13 mM tris(2-carboxyethyl)phosphine (TCEP) (Thermo Fisher Scientific, #20491) combined with 40 mM chloroacetamide (CAA) (Merck, #C0267) and incubated for 10 minutes at 95°C under agitation. Samples were acidified by adding 2.5 μL of 27% Phosphoric acid (final 2.5%, v/v) (Merck, #438081). Samples were mixed with 90% methanol (Thermo Fisher Scientific, #A456-212) and 100 mM TEAB and loaded onto S-TrapTM micro columns (≤100 μg) (ProtiFi, #C2-micro) according to manufacturer’s protocol. Proteins were digested with 2 μg trypsin (#V5111, Promega) in 20 μL of 50 mM TEAB at pH 8.0 for overnight at 37°C on the S-trap. Peptides were eluted by the sequential addition of three elution buffers containing: 50 mM TEAB (buffer 1), 0.2% formic acid (FA) (Thermo Fisher Scientific, #270480010) (buffer 2) and 50% acetonitrile (ACN) (Thermo Fisher Scientific, #A955-212) with 0.2% FA (buffer 3). Peptides were dried under reduced pressure and desalted a second time using OMIX C18 pipette (#A57003MB, Agilent Technologies). Purified peptides were completely dried and stored at −20°C until LC-MS/MS analysis. Peptides were re-dissolved in 33 µL loading solvent (0.1% TFA in H_2_O/ACN, 99.5:0.5, v/v)) of which 2 µg of the sample measured on Dropsense16 (Unchained Labs) was injected for LC-MS/MS analysis on an Ultimate 3000 Pro Flow nanoLC system in-line connected to a Q Exactive HF mass spectrometer (Thermo Fisher Scientific).

### LC-MS/MS analysis

Trapping was performed at 20 μL/min for 2 minutes in loading solvent on a 5 mm trapping column (Thermo Fisher Scientific, Pepmap, 300 μm internal diameter (I.D.), 5 μm beads). Peptides were separated on a 250 mm Aurora Ultimate, 1.7 µm C18, 75 µm inner diameter (Ionopticks) kept at a constant temperature of 45 °C. Peptides were eluted by a non-linear gradient starting at 0.5% MS solvent B (0.1% FA in ACN) and MS solvent A (0.1% FA in water). Gradient reached 33% at 70 minutes, 55% at 85 minutes and 70% at 90 minutes of MS solvent B. The gradient was followed by a 5-minute wash at 70% MS solvent B and re-equilibration with MS solvent A for 15 minutes. The mass spectrometer was operated in data-dependent mode (DDA), automatically switching between MS and MS/MS acquisition with a top 12 method. Full-scan MS spectra ranging from 375-1500 m/z with a target value of 3E6, a maximum fill time of 50 ms and a resolution at of 60,000 (at 200 m/z). Target accumulation was set at 3E6 for 60 ms in MS1. The 12 most intense ions above a threshold value of 1.3E4 were isolated in the trap with an isolation window of 1.5 Da for maximum 60 ms to a target AGC value of 1E3. Only precursor ions with a charge state equal to 2–6 were selected. Peptide match was set on “preferred” and isotopes were excluded. Dynamic exclusion time was set to 12 s. Fragmentation were performed at a normalised collision energy of 30% (NCE) with HCD fragmentation. MS/MS spectra were acquired at fixed first mass 145 m/z at a resolution of 15,000 (at 200 m/z) in the Orbitrap analyser. MS1 spectrum data type was set to profile MS2 spectrum data type was set to centroid. The polydimethylcyclosiloxane background ion at 445.12003 Da was used for internal calibration (lock mass) and QCloud was used to control instrument longitudinal performance during the project^74^.

### Data analysis for LC-MS/MS

LC-MS/MS runs of all samples were searched using the MaxQuant algorithm (version 2.2.0.0)^75^. Spectra were searched against the protein sequences in the Swiss-Prot database (www.uniprot.org), from human (UP000005640_9606) and *Shigella flexneri* (strain 301 / Serotype 2a) (UP000001006_623) (downloaded in May 2023). Additionally, the database included the protein sequence for GFP-APEX2. Mainly default settings were used. The false discovery rate was set at 1% on peptide (precursor) and protein level. Enzyme specificity was set as C-terminal to arginine and lysine, also allowing cleavage at proline bonds with a maximum of two missed cleavages. Variable modifications were set to oxidation of methionine residues, acetylation of protein Ntermini and Biotin of lysine residues. Matching between runs was turn OFF. Proteins were quantified with MaxLFQ intensities based on razor and unique peptides with a minimum of two ratio counts per protein. Further data analysis was performed with an in-house script in the R programming language by loading the proteinGroups table from MaxQuant. Briefly, proteins identified in the reverse database were removed. Protein expression matrices were prepared as follows: Only proteins with intensity in three replicates were retained for analysis. Missing intensity values were imputed by randomly sampling from a normal distribution centered around each sample’s noise level. Fold change and adjusted p-value at FDR = 0.05 were calculated with the package Limma in R^76^. GO-term enrichment analysis was performed using the Term Enrichment Service from AmiGO^77^ (https://amigo.geneontology.org/amigo) and plotted using R.

### Yeast-Two-Hybrid Screening

The Yeast-Two-Hybrid bait vector pJB79-10 encoding IpaH9.8 C337A was transformed into the Matchmaker Gold strain of *Saccharomyces cerevisiae* (Takara). A high-complexity normalised human cDNA Mate and Plate library (Takara) was screened following the protocol provided by the Matchmaker Gold Yeast-Two-Hybrid System (Takara). Positive prey clones were identified by plasmid rescue and DNA sequencing according as previously described^79^.

### Myc Fusion Protein Co-Immunoprecipitation of UFM1

10^7^ HEK293T cells were transfected with equal amounts of pRK5Myc-IpaH9.8 and pCR3.1His-UFM1, or pCR3.1His-UFM1 alone as a control. Cells were collected in PBS 24 hours post-transfection, and washed once in PBS at 4°C. Samples were resuspended in 500 μL – 1mL of lysis buffer (2% SDS, 10% glycerol, 62.5 mM Tris pH 6.8), lysed with a 21G needle or with sonication and clarified by centrifugation at 13,000 xg for 15 minutes at 4°C. Supernatants were incubated overnight with 4 volumes of cold IP Buffer (50mM Tris pH8.0, 150mM NaCl, 1% NP-40) and αMyc-Tag (71D10) antibody (Cell Signaling, #2278) at 1:1000 dilution. The next day, 1 mL of Protein A agarose beads (Sino Biological, #10600-P07E) pre-washed with IP buffer was added to the lysates and incubated for 1-3 hours. Beads were collected by centrifugation at 700 xg for 2 minutes and washed three times with 1 volume IP buffer for 10 minutes. To elute bound proteins, beads were resuspended in 2x Laemmli buffer containing 10% *β*-mercaptoethanol and boiled for 5 minutes. Supernatant was collected by centrifugation at 300 xg for 2 minutes and analysed via SDS-PAGE.

### Fluorescence polarisation binding assay

The fluorescent substrate Rhodamine-UFM1 (Rho-UFM1) was chemically synthesised as described previously^80^. Rho-UFM1, GST-IpaH9.8, and GST-ubiquitin were diluted to ‘2x’ concentrations in 25 mM Tris-HCl, 150 mM NaCl, 2 mM *β*-mercaptoethanol, 0.1 mg/mL BSA, pH 7.4. To set up the binding assay, 10 µL of 200 nM Rho-UFM1 was mixed with buffer or with 2x GST-tagged protein. The mixture was left to equilibrate in the dark at room temperature for 15 minutes, after which time it was loaded into black Greiner low-volume 384-well plates. Fluorescence polarisation values were measured on a BMG Labtech CLARIOstar plate reader equipped with 482-16 nm and 530-40 optic filters. A target polarisation value of 180 was set for free Rho-UFM1.

### Infection of HeLa cells

Hela cells were seeded in 6-well plates (Thermo Fisher Scientific) at a confluency of 1.5 x 10^5^ cells/well 2 days prior to infection. Cell cultures were infected with *S. flexneri* strains as described previously^8^. Briefly, HeLa cells were infected with *S. flexneri* by spin-inoculation at 110 xg for 10 minutes at a multiplicity of infection (MOI) of 100:1 (bacteria:cell). Then, plates were placed at 37°C and 5% CO_2_ for 30 minutes. Infected cultures were washed three times with PBS pH 7.4 and incubated with fresh DMEM containing 10% hi-FBS and 50 µg/mL gentamicin at 37°C and 5% CO_2_ for up to 5 h. Infections with *S. flexneri* mutants lacking *rfaC* (hyperinvasive) or expressing the adhesin AfaE were performed at MOI 10 in the absence of spin inoculation, unless indicated otherwise.

For *S.* Typhimurium infections, 50 µL of bacterial subculture was added to wells containing HeLa cells and incubated for 30 minutes at 37°C. Following three washes with PBS, infected cultures were incubated with fresh DMEM containing 10% FBS and 50 µg/mL gentamicin at 37°C and 5% CO_2_ for up to 5 h.

### Intracellular bacteria growth

HeLa cells seeded in 6-well plates and infected with *S. flexneri* were lysed in 1 ml cold PBS containing 0.1% Triton X-100. Serial dilutions were plated in duplicate on TCS agar containing 0.01% of Congo red for *S. flexneri* and LB agar for *S.* Typhimurium. Plates were incubated for 24 hours at 37°C and colony forming units (CFUs) enumerated. CFUs counted at 4 and 5 hours post infection were normalised to 1 hour post infection to assess intracellular growth.

### Bacteria recovery from infected cells

HeLa cells were seeded into 15 cm dishes at a confluency of 5.0 x 10^6^ and infected two days later with the hyperinvasive *S. flexneri afaI* or Δ*rfaC*Δ*ipaH9.8* at MOI 10. Extraction of bacteria from infected cells was performed as described before^14^. Cells were lysed in ice cold PBS containing 0.1% Triton X-100 at 3 hours post infection and centrifuged at 300 xg for 5 minutes at 4 °C. Supernatant containing bacteria was collected and centrifuged at 16,100 xg for 10 minutes at 4 °C. The bacterial pellet was washed once with PBS, followed by bacterial lysis in 75 μl BugBuster (Merck, #70584) including 2 mM iodoacetamide for 5 minutes at room temperature. Samples were boiled after the addition of 25 μl of Laemmli buffer 4x for 10 minutes and analysed by western blot. As control, 1 mL of bacteria in TSB was collected by centrifugation at OD (600 nm) 0.6, washed twice with PBS and lysed as described above.

### Infection of THP-1 macrophages

THP-1 cells were differentiated in 8-well µ-Slides (ibidi) at a confluency of 1 x 10^5^ cells per well. Cells were infected with *S. flexneri* strains with a multiplicity of infection (MOI) of 50. Plates were incubated for 30 minutes under standard conditions before being incubated with fresh RPMI supplemented with 10% hi-FBS and 50 µg/mL gentamycin under standard conditions for 30 minutes, followed by changing to RPMI supplemented with 10% hi-FBS containing 5 µg/mL gentamycin.

### Immunofluorescence of human cells

Bacteria, coverslips or microslides containing adherent infected or uninfected human cells were washed three times with PBS pH 7.4 and fixed 15 minutes in 4% paraformaldehyde (in PBS) at room temperature. Fixed cells were washed three times with PBS and subsequently permeabilised 5 minutes with 0.1% Triton X-100 (in PBS). Cells were then washed three times in PBS and incubated 1 hour with primary antibodies diluted in PBS supplemented with 0.1% Triton X-100 and 1% bovine serum albumin (BSA, Sigma-Aldrich, #A9418) for 1 hour at room temperature or overnight at 4 °C. Cells were then washed three times in PBS and incubated for 1 hour with secondary goat antibodies diluted in PBS supplemented with 0.1% Triton X-100 and 1% BSA, and Alexa-conjugated phalloidin where indicated. Stained bacteria cultures and coverslips were washed three times with PBS and mounted on glass slides with ProLong™ Gold Antifade Mountant with DAPI stain (Invitrogen, # P36941).

### Detection of biotinylated proteins for fluorescence microscopy in human cells

Fixed HeLa cells infected with GFP-APEX2 coated *S. flexneri* were permeabilised 5 minutes with 0.1% Triton X-100 (in PBS). Endogenous biotin was blocked with 0.05% avidin (Sigma-Aldrich, #A2667) in PBS for 15 minutes followed by 3 washes in PBS, and then 0.05% biotin (Sigma-Aldrich, #B4501) in PBS for 15 minutes followed by 3 washes in PBS. Then, proximity biotinylation was performed as described above. Biotinylated proteins were detected using Alexa Fluor-488 conjugated streptavidin and samples were further processed for immunofluorescence with anti-GFP antibodies to increase the detection of GFP-APEX2.

### Ethics statements

Animal experiments were performed according to the Animals (Scientific Procedures) Act 1986 and approved by the Home Office (PPL PP5900632).

### Zebrafish husbandry

Adult zebrafish were housed in the Biological Services Facility at LSHTM. Wildtype AB embryos and transgenic *Tg(lyzC:DsRed2)^nz^*^50^ embryos (for the expression of DsRed2 in neutrophils)^81^ were obtained by natural spawning, and larvae maintained at 28.5°C in E3 media (5 mM NaCl, 0.17 mM KCl, 0.33 mM CaCl_2_, 0.33 mM MgSO_4_). For injections, larvae were anesthetised with tricaine (160 µg/mL, Sigma-Aldrich, #A5040).

### CRISPR Cas9-mediated knockdown in zebrafish larvae

Embryos were injected at the one- to two-cell stage with 1 nL of CRISPR mixture containing 1 µg/ µL of gene-specific Guide (g) RNA and 500 µg/mL Cas9 (spCas9 V3, IDT, #1081058) (**Supplementary Table 5**). gRNA was synthesised as previously described^82^. Briefly, primers containing gene-specific guide sites and a T7 promotor were annealed to a standard scaffold primer and purified (QIAquick PCR Purification Kit, Qiagen, #28104). DNA templates were in vitro transcribed (HiScribe T7 High Yield RNA Synthesis Kit NEB, #E2040S) and purified (GeneJET RNA Cleanup and Concentration Micro Kit, ThermoFisher, #K0841).

### Gene expression analysis

Groups of 10 embryos were lysed and homogenised using a 27-gauge needle in Buffer RLT and RNA extracted with the RNeasy Mini kit (Qiagen, #74104). RNA was reverse transcribed using the QuantiTect Reverse Transcription kit (Qiagen, #205311), and quantitative (q) PCR performed on a QuantStudio Real-Time PCR system with SYBR green master mix (Applied Biosystems, #10187094). Samples were run in duplicates, and expression normalised to *ef1a1l*. Data was analysed using the 2^−ΔΔ^ Ct method. qPCR primers used are listed in **Supplementary Table 5**.

### Zebrafish injections

To prepare the bacterial inocula, subcultures of either *S. flexneri* or *S.* Typhimurium at OD (600 nm) 0.6 were centrifuged at 4,000 x*g* for 4 minutes at room temperature and washed in PBS. The bacterial pellet was then resuspended in the appropriate volume of injection buffer (4% polyvinyl-pyrrolidone [Sigma-Aldrich, #PVP40] in PBS and 0.5% phenol red [Sigma-Aldrich, #114537] to obtain an infection dose of 10,000 CFU/nL for *S. flexneri* or 1,500 CFU for *S.* Typhimurium. For injection, 1 nL of bacterial suspension was microinjected into either the tail musculature (for immunofluorescence experiments) or caudal vein (for survival assays) of 3 days post fertilisation larvae.

### Wholemount zebrafish immunostaining

Zebrafish larvae were fixed at 2 hours post infection in 4% paraformaldehyde with 0.4% triton X-100 and incubated at 4°C overnight. Larvae were then washed three times for five minutes in 0.4% Triton X-100 in PBS, followed by one 20-minute wash in 1% Triton X-100 in PBS. Samples were then blocked in blocking buffer (10% hi-FBS, 1% DMSO, 0.1% Tween-20 in PBS) for one hour at room temperature. Primary antibodies were prepared in blocking buffer and larvae were incubated overnight at 4°C with gentle rocking. Larvae were washed four times for 15 minutes with 0.1% Tween-20 in PBS, before incubation overnight at 4°C with secondary antibodies prepared in blocking buffer with gentle rocking. Larvae were washed as described and then cleared in a glycerol series of 25, 50 and 70%, allowing the larvae to sink to the bottom of the tube between each glycerol concentration. Larvae were mounted on 35-mm glass-bottom dishes (#P35G-1.5-14-C; MatTek) in a minimal volume of 70% glycerol for subsequent imaging.

### Detection of biotinylated proteins for fluorescence microscopy in zebrafish larvae

Fixed zebrafish larvae infected with GFP-APEX2 coated *S. flexneri* were permeabilised with 0.4% triton X-100 and incubated at 4°C overnight. Larvae were then washed three times for five minutes in 0.4% Triton X-100 in PBS, followed by one 20-minute wash in 1% Triton X-100 in PBS. Endogenous biotin was blocked with 0.05% avidin in blocking buffer (10% hi-FBS, 1% DMSO, 0.1% Tween-20 in PBS) for one hour at room temperature, washed four times for 15 minutes, incubated with 0.05% biotin in blocking buffer for 1 hour at room temperature and washed four times for 15 minutes. Then, proximity biotinylation was performed as described above. Biotinylated proteins were detected using Alexa Fluor-488 conjugated streptavidin and samples were further processed for immunofluorescence with anti-GFP antibodies to increase the detection of GFP-APEX2.

### Bacterial recovery from zebrafish larvae

At the required timepoint, 5 larvae per group were collected for CFU recovery. Larvae were euthanised by anaesthetic overdose (tricaine) and individually homogenised with a 27-gauge needle in 150 µL of PBS. Homogenates were serially diluted and plated on TSA plates supplemented with 0.01% Congo red, or LB agar plates. Plates were incubated for 24 hours at 37°C and CFU enumerated.

### Flow cytometry

5 x 10^4^ individual bacterial cells were analysed using flow cytometry with an LSRII flow cytometer (BD Biosciences). The data was analysed using FlowJo software, version 10.7.1. The bacterial fluorescence was normalised to the median fluorescence of the control condition (no addition of GFP-APEX2) for each biological replicate. The normalised values, median and interquartile range was plotted.

### Secretion assay

Secretion of T3SS effectors was performed as previously described^83^. Briefly, bacteria were grown overnight, subcultured and grown until OD (600nm) of 0.4–0.5 at 37°C. Cultures were then incubated for 3 hours in the presence or absence of Congo red to induce Type 3 secretion. Secreted proteins were collected from culture supernatants, precipitated using trichloroacetic acid (Sigma-Aldrich, #T0699), and then analysed using SDS-PAGE and Coomassie Brilliant Blue R-250 (Bio-Rad, #1610436) staining.

### SDS-PAGE

Cells were lysed in ice-cold RIPA buffer and then incubated 1:1 with Laemmli buffer (2x) containing 10% (v/v) *β*-mercaptoethanol at 95°C for 10 minutes. Proteins were resolved in 12% or 4–20% Mini-PROTEAN TGX precast protein gels (Biorad, #4561046EDU, #4561096EDU).

### Western blotting

Protein gels were transferred to 0.2 μm polyvinylidene difluoride membranes (PVDF, Biorad, #1704156). PVDF membranes were incubated in blocking buffer (TBST buffer containing 3% of BSA) for 1 hour. PVDF membranes were incubated with the primary antibodies diluted in blocking buffer for 1 hour at room temperature or overnight at 4 °C. PVDF membranes were washed 3× 5–7 minutes in PBS at room temperature and incubated with secondary goat HRP-conjugated antibodies in blocking buffer for 1 hour at room temperature. Milk was replaced by 3% BSA in the blocking buffer for the detection of biotinylated proteins with Streptavidin-HRP or detection of UFM1 with antibodies. Membranes were developed using Pierce^TM^ ECL plus western blotting substrate (Thermo Scientific, #32132)

### Microscopy

Fluorescence microscopy was performed using a 63×/1.4 C-Plan Apo oil immersion lens or a 40x/1.2 C-Apo water immersion lens on a Zeiss LSM 880 confocal microscope driven by ZEN Black software (v2.3). Microscopy images were obtained using z-stack image series taking 8–16 slices (HeLa cells) or 40-120 slices (zebrafish larvae). When using the airyscan acquisition mode, confocal images were processed using airyscan processing (Weiner filter) using “Auto Filter” and “3D Processing” options. Alternatively, and for the purpose of quantification of host factors to bacteria, fluorescence microscopy of infected HeLa cells was performed on an AxioObserver Z1 fluorescence microscope driven by ZEN Blue v2.3 software (Carl Zeiss). Whole larvae images were acquired using stereo fluorescent microscope Leica M205FA (Leica, Germany). Image files were processed using ImageJ/Fiji software for representation.

### Statistics

All graphs were plotted, and statistical analysis were performed using Prism. n.s: non-significant, *: p-value < 0.05, **: p-value < 0.01, ***: p-value < 0.001.

## Tables

**Supplementary Table 1.** Mass spectrometry protein identification and quantification of *S. flexneri* WT or Δ*mixE* proxisomes during infection of HeLa cells. Proteins were identified with MaxQuant at 1% FDR to both peptides and protein level.

**Supplementary Table 2.**
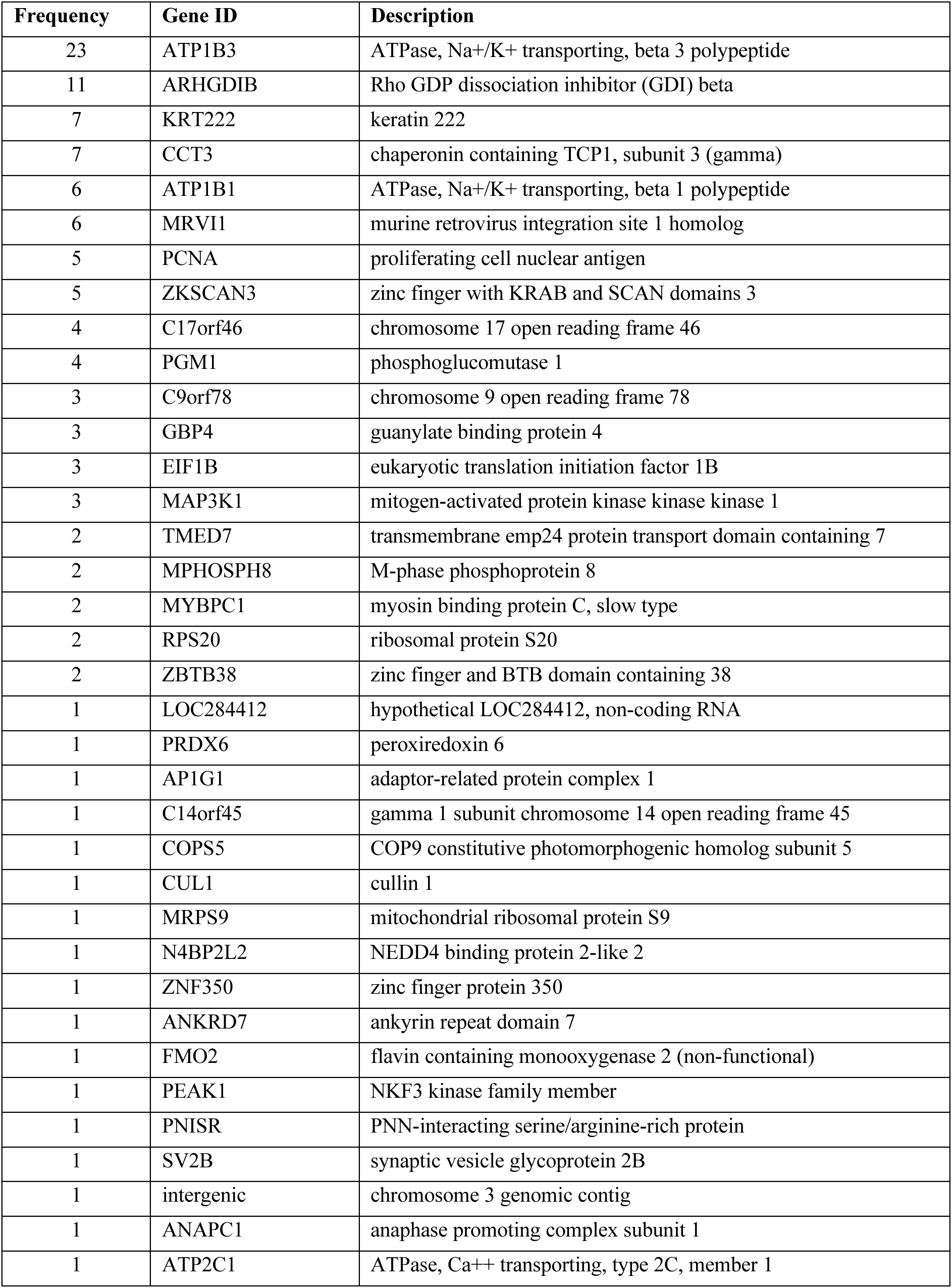
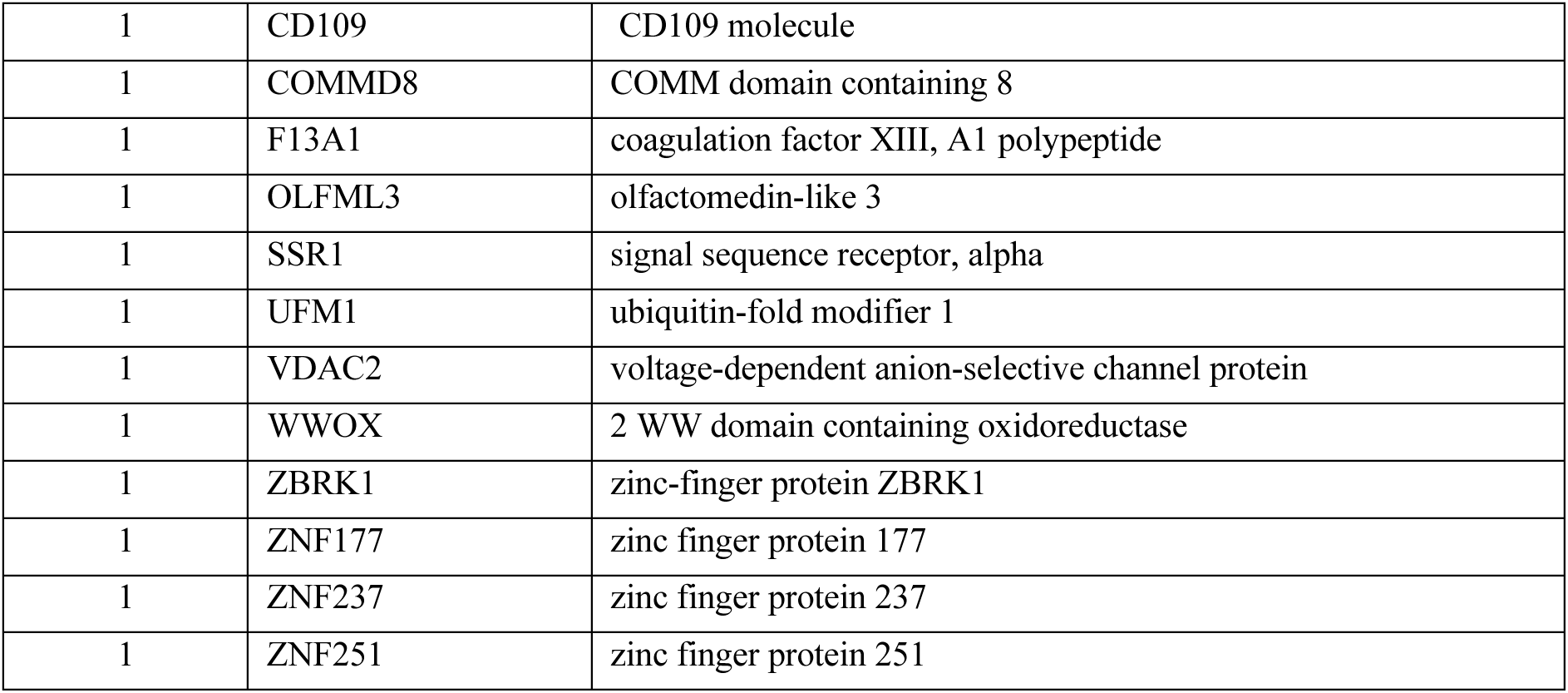
Top 50 mammalian proteins shown to interact with IpaH9.8 by Yeast-Two-Hybrid protein-protein interaction screen, ordered by the number of clones detected.

| Frequency | Gene ID | Description |
| --- | --- | --- |
| 23 | ATP1B3 | ATPase, Na<sup>+</sup>/K<sup>+</sup> transporting, beta 3 polypeptide |
| 11 | ARHGDIB | Rho GDP dissociation inhibitor (GDI) beta |
| 7 | KRT222 | keratin 222 |
| 7 | CCT3 | chaperonin containing TCP1, subunit 3 (gamma) |
| 6 | ATP1B1 | ATPase, Na<sup>+</sup>/K<sup>+</sup> transporting, beta 1 polypeptide |
| 6 | MRVI1 | murine retrovirus integration site 1 homolog |
| 5 | PCNA | proliferating cell nuclear antigen |
| 5 | ZKSCAN3 | zinc finger with KRAB and SCAN domains 3 |
| 4 | C17orf46 | chromosome 17 open reading frame 46 |
| 4 | PGM1 | phosphoglucomutase 1 |
| 3 | C9orf78 | chromosome 9 open reading frame 78 |
| 3 | GBP4 | guanylate binding protein 4 |
| 3 | EIF1B | eukaryotic translation initiation factor 1B |
| 3 | MAP3K1 | mitogen-activated protein kinase kinase kinase 1 |
| 2 | TMED7 | transmembrane emp24 protein transport domain containing 7 |
| 2 | MPHOSPH8 | M-phase phosphoprotein 8 |
| 2 | MYBPC1 | myosin binding protein C, slow type |
| 2 | RPS20 | ribosomal protein S20 |
| 2 | ZBTB38 | zinc finger and BTB domain containing 38 |
| 1 | LOC284412 | hypothetical LOC284412, non-coding RNA |
| 1 | PRDX6 | peroxiredoxin 6 |
| 1 | AP1G1 | adaptor-related protein complex 1 |
| 1 | C14orf45 | gamma 1 subunit chromosome 14 open reading frame 45 |
| 1 | COPS5 | COP9 constitutive photomorphogenic homolog subunit 5 |
| 1 | CUL1 | cullin 1 |
| 1 | MRPS9 | mitochondrial ribosomal protein S9 |
| 1 | N4BP2L2 | NEDD4 binding protein 2-like 2 |
| 1 | ZNF350 | zinc finger protein 350 |
| 1 | ANKRD7 | ankyrin repeat domain 7 |
| 1 | FMO2 | flavin containing monooxygenase 2 (non-functional) |
| 1 | PEAK1 | NKF3 kinase family member |
| 1 | PNISR | PNN-interacting serine/arginine-rich protein |
| 1 | SV2B | synaptic vesicle glycoprotein 2B |
| 1 | intergenic | chromosome 3 genomic contig |
| 1 | ANAPC1 | anaphase promoting complex subunit 1 |
| 1 | ATP2C1 | ATPase, Ca<sup>++</sup> transporting, type 2C, member 1 |
| 1 | CD109 | CD109 molecule |
| 1 | COMMD8 | COMM domain containing 8 |
| 1 | F13A1 | coagulation factor XIII, A1 polypeptide |
| 1 | OLFML3 | olfactomedin-like 3 |
| 1 | SSR1 | signal sequence receptor, alpha |
| 1 | UFM1 | ubiquitin-fold modifier 1 |
| 1 | VDAC2 | voltage-dependent anion-selective channel protein |
| 1 | WWOX | 2 WW domain containing oxidoreductase |
| 1 | ZBRK1 | zinc-finger protein ZBRK1 |
| 1 | ZNF177 | zinc finger protein 177 |
| 1 | ZNF237 | zinc finger protein 237 |
| 1 | ZNF251 | zinc finger protein 251 |

**Supplementary Table 3.**
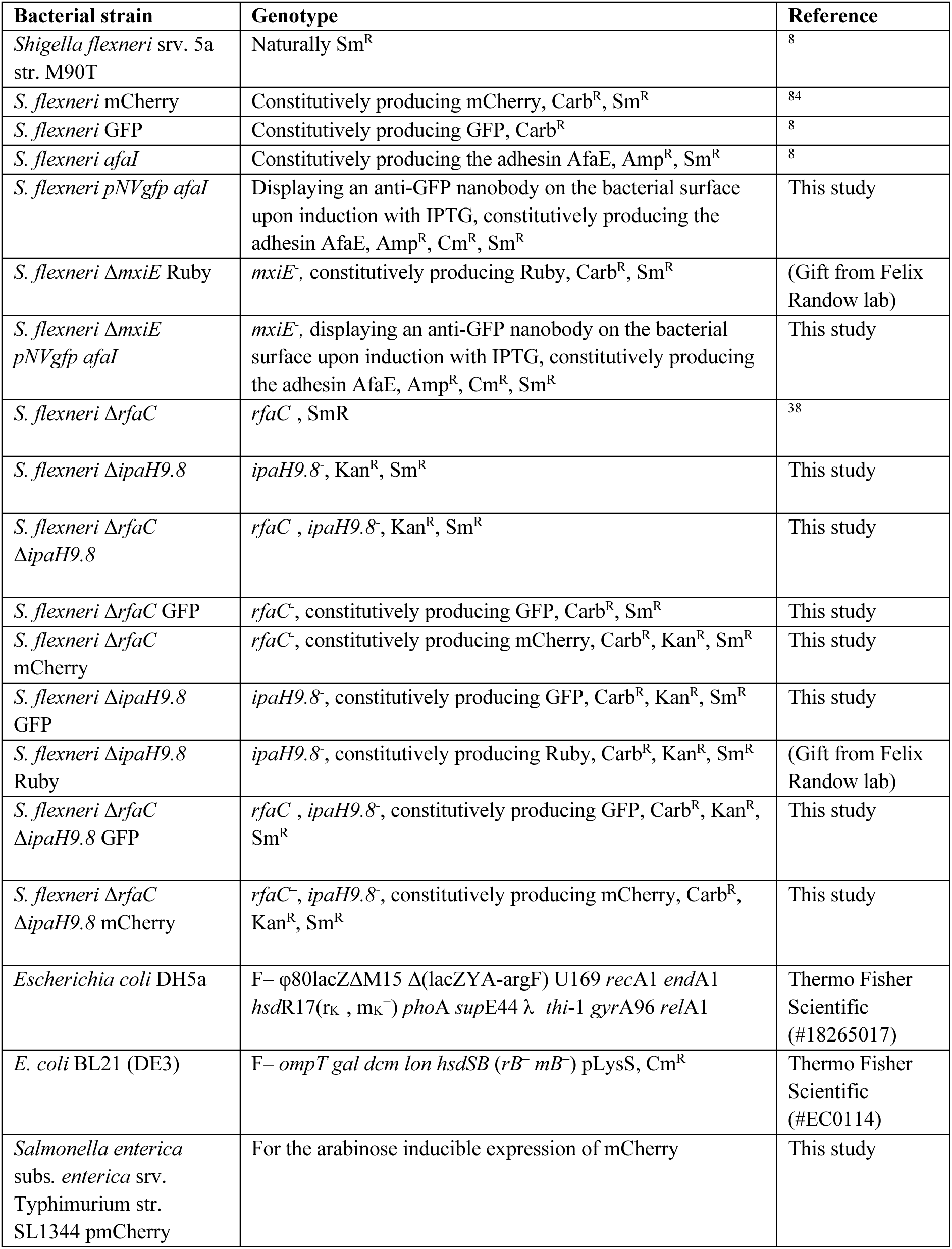
Bacterial strains used in this study.

**Supplementary Table 4.**
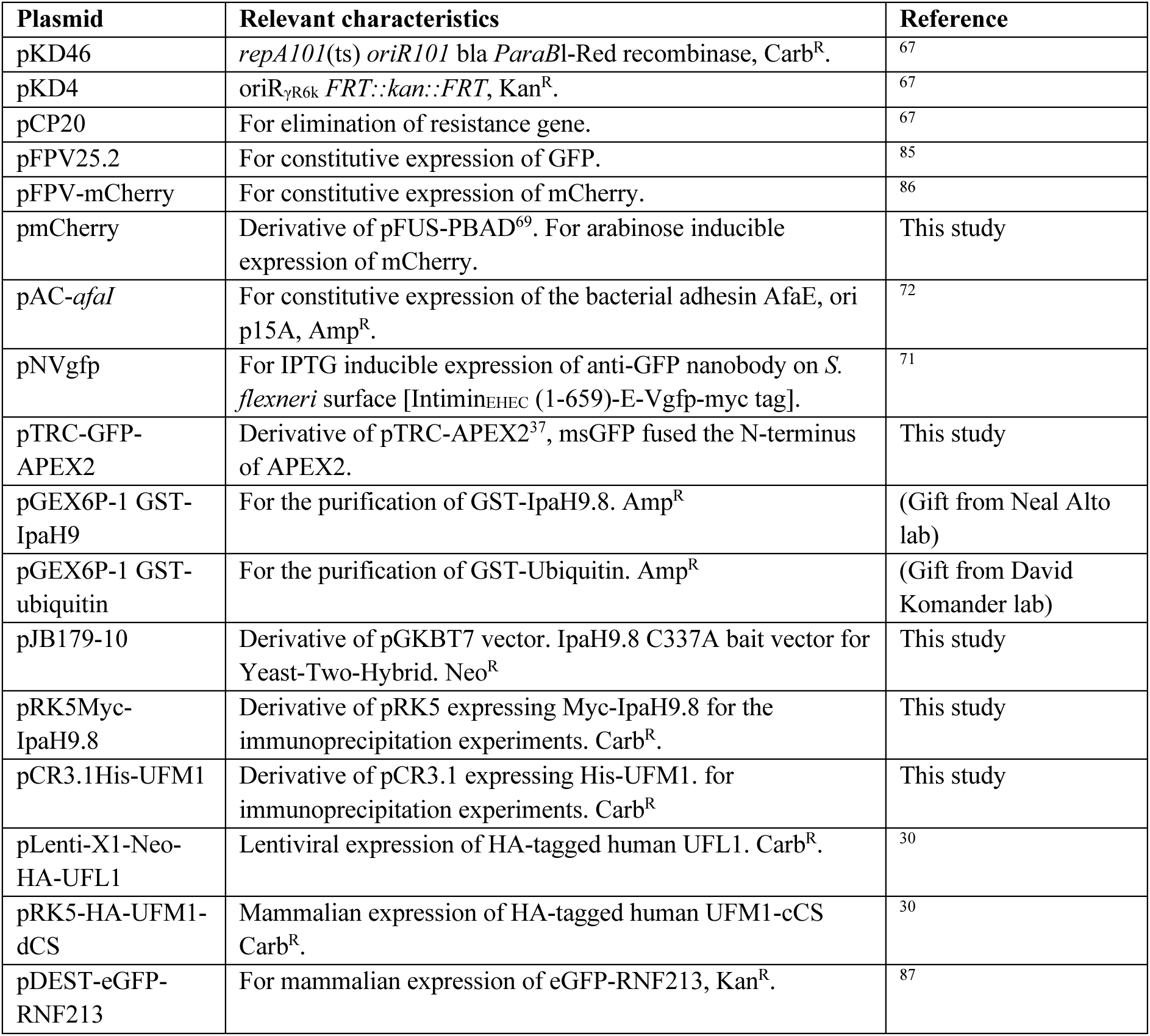
Plasmids used in this study.

**Supplementary Table 5.**
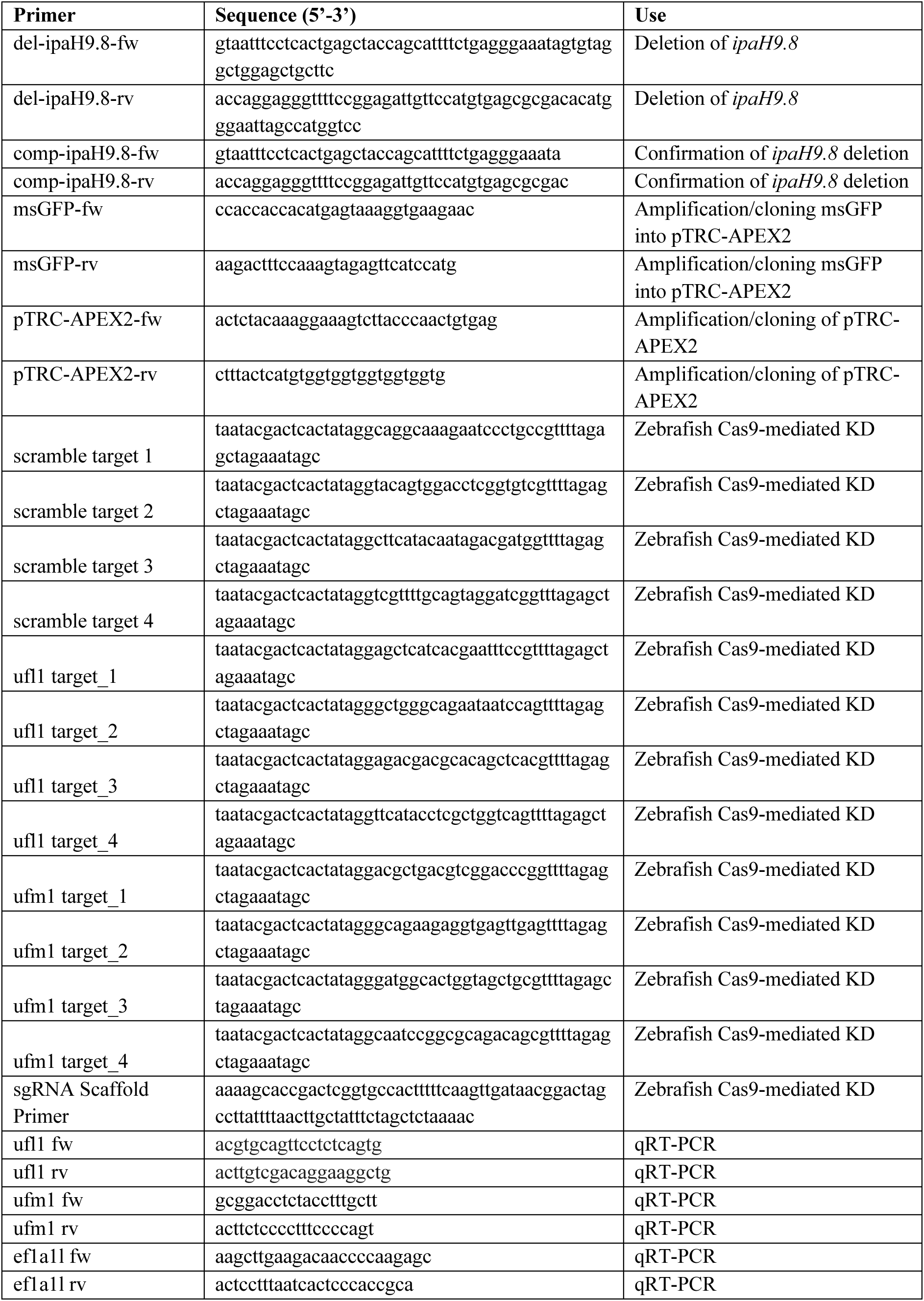
Primers used in this study.

## Extended Data Figures

**Extended Data Fig. 1. Functionalisation of *S. flexneri* with GFP-APEX2 and proximity biotinylation of bacterial surface in vitro. a,** Diagram representing the GFP-APEX2 construct for *E. coli* expression and subsequent purification. **b,** Purification of GFP-APEX2. Coomassie stained SDS-PAGE showing protein ladder for molecular weight (MW), clarified extract (CE), flowthrough (FT), elution fractions (E) and purified protein (PP). Fractions E2 to E5 were pulled together. Resulting GFP-APEX2 protein has a predicted molecular weight of 55.69 KDa. **c,** Diagram showing the experimental method. First, the expression is anti-GFP nanobody on *S. flexneri* surface is induced with IPTG. Then, bacteria are functionalised in vitro with GFP-APEX2. Finally, the proximity biotinylation reaction happens after addition of BP and H_2_O_2_. **d,** Representative airyscan confocal images showing biotinylation at the vicinity of functionalised *S. flexneri* in vitro, in the presence or absence of IPTG and H_2_O_2_. SAV stands for streptavidin. Scale bar, 1 µm. **e,** Quantification of GFP-APEX2 coating and protein biotinylation at the surface of *S. flexneri* in the presence or absence of GFP-APEX2, BP and H_2_O_2_, measured by flow cytometry. More than 60,000 bacteria from a total of 3 independent experiments were analysed per condition. The results are represented as median ± interquartile range. Kruskal-Wallis test and Dunn’s multiple comparisons test.

**Extended Data Fig. 2. Functionalised *S. flexneri* infects HeLa cells. a-b,** Representative confocal microscopy images of HeLa cells infected with GFP-APEX2 coated *S. flexneri* forming actin tails and entrapped in septin cages, respectively. **c,** In vitro secretion assay of *S. flexneri* WT and Δ*mxiE* displaying surface nanobodies and producing the AfaE adhesin indicate a functional T3SS upon Congo red stimulation. **d,** *S. flexneri* WT and Δ*mxiE* displaying surface nanobodies and producing the AfaE adhesin have similar HeLa invasion rates. The results are represented as mean ± SD.

**Extended Data Fig. 3. *S. flexneri* proxisome leads to identification of novel host factors recruited to bacteria. a,** Principal component analysis showing different proxisomes are identified during infection of *S. flexneri* WT and Δ*mxiE.* **b,** Volcano plot of proteins identified. **c,** GO terms for biological process enriched in the proxisome of *S. flexneri* during infection. **d,** STRING diagram displaying the E3 ligases and deubiquitylases identified. Edges represent shared physical complex; strength of the edge represents confidence.

**Extended Data Fig. 4. UFL1 and UFM1 are recruited to *S flexneri*.** Representative confocal images of HeLa cells expressing HA-UFL1 or HA-UFM1-dCs and infected with *S. flexneri afaI.* Arrows show localisation of exogenous proteins to intracellular bacteria. Scale bar, 10 µm.

**Extended Data Fig. 5. Deletion of ipaH9.8 in *S. flexneri* and recruitment UFL1 and UFM1 recruitment to *S. flexneri* and *S.* Typhimurium in HeLa cells. a**, Deletion of *ipaH9.8* in *S. flexneri* WT and Δ*rfaC*. Diagram indicating the primer annealing regions for the insertion of a Kanamycin resistance cassette, prior (top) and after (bottom) insertion. **b-c**, PCR showing the insertion of the Kanamycin resistance cassette in *S. flexneri* WT and Δ*rfaC.* **d,** Percentage of *S. flexneri* WT (n=1311, n=837), Δ*ipaH9.8* (n=463, n=576), Δ*rfaC* (n=630, n=721), and double Δ*rfaC*Δ*ipaH9.8* (n=553, n=473) mutants colocalising with UFL1 at 3 and 5 hours post infection (h.p.i), respectively. **e,** Percentage of *S. flexneri* WT (n=882, n=566), Δ*ipaH9.8* (n=370, n=531), Δ*rfaC* (n=643, n=851), and double Δ*rfaC*Δ*ipaH9.8* (n=526, n=650) mutants colocalising with UFM1 at 3 and 5 hours post infection, respectively. The results are represented as mean ± SD. Two-way ANOVA and Tukey’s multiple comparison test. **f,** Representative deconvolved widefield image showing HeLa cell expressing GFP-RNF213 infected with *S.* Typhimurium and immunostained for UFL1. Green, magenta and white arrows show bacteria recruited with GFP-RNF213 only, UFL1 only, or both, respectively. Scale bar, 5 µm.

**Extended Data Fig. 6. IpaH9.8 immunoprecipitation leads to UFM1 species enrichment.** HEK293T cells were transfected with plasmids that express His-UFM1 and/or Myc-IpaH9.8 and harvested. Unbound (UB), wash (W1, W2, W3), and eluted (ELU) fractions were analysed via SDS-PAGE followed by Coomassie stain or immunoblot. His-UFM1 was co-purified with Myc-IpaH9.8 using protein anti-myc antibody agarose beads, versus a His-UFM1-only control. UFM1 species were detected with anti-UFM1 antibodies (BostonBiochem).

**Extended Data Fig. 7. UFL1 and UFM1 are recruited to intracellular *S. flexneri* in THP-1 macrophages and in zebrafish larvae. a-b,** Representative confocal image of THP-1 macrophages infected with *S. flexneri* WT, Δ*rfaC*, and Δ*rfaC*Δ*ipaH9.8*. Arrows show localisation of UFL1 and UFM1, respectively, to intracellular bacteria. Scale bar, 10 µm. **c-d,** Representative confocal image of zebrafish larvae infected at the tail musculature with *S. flexneri* WT, Δ*rfaC*, and Δ*rfaC*Δ*ipaH9.8* at 2 hours post infection and stained against UFL1 or UFM1. Scale bar, 20 µm and 5 µm for the inset. Arrows show localisation of UFL1 or UFM1 to bacteria. Yellow arrows show bacteria that have partially or completely lost their mCherry fluorescence.

**Extended Data Fig. 8. Knockdown of *ufm1* in HeLa cells. a,** Western blot showing reduced UFM1 levels after depletion by siRNA in HeLa cells, using anti-UFM1 antibodies (Proteintech). **b,** Quantification of UFM1 upon siRNA depletion detected by Western blot. Student’s t-test. **c,** Representative microscopy images of HeLa cells infected with S*. flexneri* Δ*rfaC*Δ*ipaH9.8* immunostained against UFM1, under control conditions or after siRNA depletion, confirming antibody specificity. **d-f,** Invasion of *S. flexneri*, *S. flexneri* Δ*rfaC*Δ*ipaH9.8* and *S.* Typhimurium in HeLa cells. Invasion is calculated as the ratio of CFUs recovered at 1 hour post infection (internalised bacteria) to the CFUs of the inoculum. Student’s t-test. **g-i,** Intracellular growth of *S. flexneri*, *S. flexneri* Δ*rfaC*Δ*ipaH9.8* and *S.* Typhimurium in HeLa cells, normalised to 1 hour post infection. The results are represented as mean ± SD.

**Extended Data Fig. 9. Characterisation of *ufl1* and *ufm1* CRISPR/Cas9 knockdown zebrafish larvae. a-b,** Quantification of *ufm1* and *ufl1* expression in zebrafish larvae upon CRISPR/Cas9 knockdown by qRT-PCR. Values are normalised to the median of the control (scramble) condition. The results are represented as mean ± SD. Student’s t-test. **c-d,** Representative microscopy images of zebrafish larvae infected with *S. flexneri* at the tail musculature at 2 hours post infection and immunostained against UFM1 and UFL1, respectively, upon *ufm1* and *ufl1* depletion or control conditions, confirming specificity of antibodies. Scale bar, 20 µm. **e,** Representative images of CRISPR Cas9 generated zebrafish larvae with scramble, *ufl1* or *ufm1* gRNAs at 2-, 3- and 5-days post fertilisation (d.p.f.).

**Extended Data Fig. 10. Proposed model for intracellular bacteria UFMylation.** UFL1 is recruited to the *S. flexneri* surface, where it mediates the decoration of the bacteria with UFM1 to target them for degradation in a mechanism independent of autophagy. UFL1 recruitment is hampered by bacterial LPS, and the Type 3 secreted effector IpaH9.8 binds to UFM1 to prevent its deposition on *S. flexneri*.

## Additional information

**Supplementary Video 1. Proximity biotinylation in zebrafish larvae.** 3D reconstruction of a confocal microscopy z-stack showing a *Tg(lyzC:DsRed2)^nz50^* zebrafish larva with fluorescently labelled neutrophils infected at the tail musculature with S. *flexneri* WT functionalised for proximity biotinylation. GFP-APEX2 coated bacteria (green) have been phagocytosed by neutrophils (red). Biotinylated proteins (magenta) appear around phagocytosed bacteria. The 3D grid overlay corresponds to 30 µm intervals.

## References

1. Khalil, I. A. et al. Morbidity and mortality due to *Shigella* and enterotoxigenic *Escherichia coli* diarrhoea: the Global Burden of Disease Study 1990-2016. Lancet Infect Dis 18, 1229– 1240 (2018).

2. Bajunaid, W. et al. The T3SS of *Shigella*: expression, structure, function, and role in vacuole escape. Microorganisms Vol. 8, Page 1933 8, 1933 (2020).

3. Pinaud, L. et al. Identification of novel substrates of *Shigella* T3SA through analysis of its virulence plasmid-encoded secretome. PLoS One 12, e0186920 (2017).

4. Ray, K. et al. Tracking the dynamic interplay between bacterial and host factors during pathogen-induced vacuole rupture in real time. Cell Microbiol 12, 545–556 (2010).

5. Bernardini, M. L., Mounier, J., D’Hauteville, H., Coquis-Rondon, M. & Sansonetti, P. J. Identification of *icsA*, a plasmid locus of *Shigella flexneri* that governs bacterial intra- and intercellular spread through interaction with F-actin. Proc Natl Acad Sci U S A 86, 3867–3871 (1989).

6. Egile, C. et al. Activation of the Cdc42 effector N-Wasp by the *Shigella flexneri* Icsa protein promotes actin nucleation by Arp2/3 complex and bacterial actin-based motility. J Cell Biol 146, 1319–1332 (1999).

7. Randow, F., MacMicking, J. D. & James, L. C. Cellular self-defense: How cell-autonomous immunity protects against pathogens. Science (1979) 340, 701–706 (2013).

8. Mostowy, S. et al. Entrapment of intracytosolic bacteria by septin cage-like structures. Cell Host Microbe 8, 433–444 (2010).

9. Krokowski, S. et al. Septins recognize and entrap dividing bacterial cells for delivery to lysosomes. Cell Host Microbe 24, 866–874.e4 (2018).

10. Wandel, M. P. et al. GBPs inhibit motility of *Shigella flexneri* but are targeted for degradation by the bacterial ubiquitin ligase IpaH9.8. Cell Host Microbe 22, 507–518.e5 (2017).

11. Li, P. et al. Ubiquitination and degradation of GBPs by a *Shigella* effector to suppress host defence. Nature 551, 378–383 (2017).

12. Kutsch, M. et al. Direct binding of polymeric GBP1 to LPS disrupts bacterial cell envelope functions. EMBO J 39(13):e104926 (2020).

13. Gaudet, R. G. et al. A human apolipoprotein L with detergent-like activity kills intracellular pathogens. Science 373(6552):eabf8113 (2021).

14. Otten, E. G. et al. Ubiquitylation of lipopolysaccharide by RNF213 during bacterial infection. Nature 594, 111–116 (2021).

15. Ogawa, M. et al. Escape of intracellular *Shigella* from autophagy. Science 307, 727–731 (2005).

16. Baxt, L. A. & Goldberg, M. B. Host and bacterial proteins that repress recruitment of LC3 to *Shigella* early during infection. PLoS One 9(4):e94653 (2014).

17. Campbell-Valois, F. X., Sachse, M., Sansonetti, P. J. & Parsot, C. Escape of actively secreting *Shigella flexneri* from ATG8/LC3-Positive vacuoles formed during cell-to-cell spread is facilitated by IcsB and VirA. mBio 6, 1–11 (2015).

18. Naydenova, K., et al. *Shigella flexneri* evades LPS ubiquitylation through IpaH1.4-mediated degradation of RNF213. Nat Struct Mol Biol (2025) doi:10.1038/s41594-025-01530-8.

19. Zhou, X., et al. *Shigella* effector IpaH1.4 subverts host E3 ligase RNF213 to evade antibacterial immunity. Nat Commun 16(1):3099 (2025).

20. Saavedra-Sanchez, L. et al. The *Shigella flexneri* effector IpaH1.4 facilitates RNF213 degradation and protects cytosolic bacteria against interferon-induced ubiquitylation. Elife 13:RP102714 (2024).

21. Cappadocia, L. & Lima, C. D. Ubiquitin-like protein conjugation: structures, chemistry, and mechanism. Chem Rev 118, 889 (2018).

22. Spinnenhirn, V. et al. The ubiquitin-like modifier FAT10 decorates autophagy-targeted *Salmonella* and contributes to *Salmonella* resistance in mice. J Cell Sci 127, 4883–4893 (2014).

23. Radoshevich L et al. ISG15 counteracts *Listeria monocytogenes* infection. Elife 4, (2015).

24. Jubelin, G. et al. Pathogenic bacteria target NEDD8-conjugated cullins to hijack host-cell signaling pathways. PLoS Pathog 6, e1001128 (2010).

25. McCormack, R. M., Lyapichev, K., Olsson, M. L., Podack, E. R. & Munson, G. P. Enteric pathogens deploy cell cycle inhibiting factors to block the bactericidal activity of Perforin-2. Elife 4:e06505 (2015).

26. Sidik, S. M., Salsman, J., Dellaire, G. & Rohde, J. R. *Shigella* infection interferes with SUMOylation and increases PML-NB number. PLoS One 10, e0122585 (2015).

27. Lapaquette, P., et al. *Shigella* entry unveils a calcium/calpain-dependent mechanism for inhibiting sumoylation. Elife 6:e27444, (2017).

28. Picchianti, L. et al. Shuffled ATG8 interacting motifs form an ancestral bridge between UFMylation and autophagy. EMBO J 42(10):e112053 (2023).

29. Wang, L. et al. UFMylation of RPL26 links translocation-associated quality control to endoplasmic reticulum protein homeostasis. Cell Res 30:1 30, 5–20 (2019).

30. Liang, J. R. et al. A genome-wide ER-phagy screen highlights key roles of mitochondrial metabolism and ER-resident UFMylation. Cell 180, 1160–1177.e20 (2020).

31. Qin, B. et al. UFL1 promotes histone H4 ufmylation and ATM activation. Nat Commun 10:1 10, 1–13 (2019).

32. Lee, L. et al. UFMylation of MRE11 is essential for telomere length maintenance and hematopoietic stem cell survival. Sci Adv 7, 7371–7395 (2021).

33. Eck, F. et al. ACSL3 is a novel GABARAPL2 interactor that links ufmylation and lipid droplet biogenesis. J Cell Sci 133(18):jcs243477 (2020).

34. Dejesus, R. et al. Functional CRISPR screening identifies the ufmylation pathway as a regulator of SQSTM1/p62. Elife 5:e17290 (2016).

35. Balce, D. R. et al. UFMylation inhibits the proinflammatory capacity of interferon-γ– activated macrophages. Proc Natl Acad Sci U S A 118, e2011763118 (2021).

36. Parsot, C. *Shigella* type III secretion effectors: how, where, when, for what purposes? Curr Opin Microbiol 12, 110–116 (2009).

37. Lam, S. S. et al. Directed evolution of APEX2 for electron microscopy and proximity labeling. Nat Methods 12, 51–54 (2014).

38. Lobato-Márquez, D. et al. Mechanistic insight into bacterial entrapment by septin cage reconstitution. Nat Commun 12:1 12, 1–14 (2021).

39. Birmingham, C. L., Smith, A. C., Bakowski, M. A., Yoshimori, T. & Brumell, J. H. Autophagy controls *Salmonella* infection in response to damage to the *Salmonella*-containing vacuole. J Biol Chem 281, 11374–11383 (2006).

40. Rohde, J. R., Breitkreutz, A., Chenal, A., Sansonetti, P. J. & Parsot, C. Type III secretion effectors of the IpaH family are E3 ubiquitin ligases. Cell Host Microbe 1, 77–83 (2007).

41. Huibregtse, J. & Rohde, J. R. Hell’s BELs: bacterial E3 ligases that exploit the eukaryotic ubiquitin machinery. PLoS Pathog 10, e1004255 (2014).

42. Ashida, H. & Sasakawa, C. *Shigella* IpaH family effectors as a versatile model for studying pathogenic bacteria. Front Cell Infect Microbiol 5, 172416 (2016).

43. Mousnier, A. et al. A new method to determine in vivo interactomes reveals binding of the *Legionella pneumophila* effector PieE to multiple rab GTPases. mBio 5(4):e01148–14 (2014).

44. D’Costa, V. M., et al. BioID screen of *Salmonella* type 3 secreted effectors reveals host factors involved in vacuole positioning and stability during infection. Nat Microbiol 4, 2511–2522 (2019).

45. Zoued, A. et al. Proteomic analysis of the host–pathogen interface in experimental cholera. Nat Chem Biol 2021 17:11 17, 1199–1208 (2021).

46. Howe, D. & Heinzen, R. A. Fractionation of the *Coxiella burnetii* parasitophorous vacuole. Methods Mol Biol 445, 389–406 (2008).

47. Lee, B. Y., et al. The *Mycobacterium bovis* bacille Calmette-Guérin phagosome proteome. Mol Cell Proteomics 9, 32–53 (2010).

48. He, Y., Li, W., Liao, G. & Xie, J. *Mycobacterium tuberculosis*-specific phagosome proteome and underlying signaling pathways. J Proteome Res 11, 2635–2643 (2012).

49. Hoffmann, C., Finsel, I. & Hilbi, H. Pathogen vacuole purification from *Legionella-*infected amoeba and macrophages. Methods Mol Biol 954, 309–321 (2013).

50. Cheng, Y., et al. Proteomic analysis of the *Ehrlichia chaffeensis* phagosome in cultured DH82 cells. PLoS One 9, e88461 (2014).

51. Murray, G. L., Attridge, S. R. & Morona, R. Altering the length of the lipopolysaccharide O-antigen has an impact on the interaction of *Salmonella enterica* serovar Typhimurium with macrophages and complement. J Bacteriol 188, 2735 (2006).

52. Domínguez-Medina, C. C. et al. Outer membrane protein size and LPS O-antigen define protective antibody targeting to the *Salmonella* surface. Nat Commun 11(1):851 (2020).

53. Shi, J. et al. Inflammatory caspases are innate immune receptors for intracellular LPS. Nature 514, 187–192 (2014).

54. Gaudet, R. G., et al. Innate recognition of intracellular bacterial growth is driven by the TIFA-dependent cytosolic surveillance pathway. Cell Rep 19, 1418–1430 (2017).

55. Milivojevic, M. et al. ALPK1 controls TIFA/TRAF6-dependent innate immunity against heptose-1,7-bisphosphate of gram-negative bacteria. PLoS Pathog 13, e1006224 (2017).

56. Morona, R., Daniels, C. & Van Den Bosch, L. Genetic modulation of *Shigella flexneri* 2a lipopolysaccharide O antigen modal chain length reveals that it has been optimized for virulence. Microbiology (N Y) 149, 925–939 (2003).

57. Paciello, I. et al. Intracellular *Shigella* remodels its LPS to dampen the innate immune recognition and evade inflammasome activation. Proc Natl Acad Sci U S A 110, E4345 (2013).

58. Okuda, J., et al. *Shigella* effector IpaH9.8 binds to a splicing factor U2AF35 to modulate host immune responses. Biochem Biophys Res Commun 333, 531–539 (2005).

59. Ashida, H. et al. A bacterial E3 ubiquitin ligase IpaH9.8 targets NEMO/IKKγ to dampen the host NF-κB-mediated inflammatory response. Nat Cell Biol 12, 66–73 (2010).

60. Li, C. et al. UFL1 alleviates lipopolysaccharide-induced cell damage and inflammation via regulation of the TLR4/NF-κB pathway in bovine mammary epithelial cells. Oxid Med Cell Longev 2019:6505373 (2019).

61. Snider, D. L., Park, M., Murphy, K. A., Beachboard, D. C. & Horner, S. M. Signaling from the RNA sensor RIG-I is regulated by ufmylation. Proc Natl Acad Sci U S A 119(15):e2119531119 (2022).

62. Tao, Y. et al. UFL1 promotes antiviral immune response by maintaining STING stability independent of UFMylation. Cell Death Differ 30:1 30, 16–26 (2022).

63. Garelis, N. E., Luteijn, R. D., Raulet, D. H. & Cox, J. S. UFMylation suppresses Type I IFN signaling during *M. tuberculosis* infection of human macrophages. bioRxiv 2024.08.07.607094 (2024)

64. Mavris, M. et al. Regulation of transcription by the activity of the *Shigella flexneri* type III secretion apparatus. Mol Microbiol 43, 1543–1553 (2002).

65. Runwal, G. et al. LC3-positive structures are prominent in autophagy-deficient cells. Sci Rep 2019 9:1 9, 1–14 (2019).

66. Chanput, W., Mes, J. J. & Wichers, H. J. THP-1 cell line: An in vitro cell model for immune modulation approach. Int Immunopharmacol 23, 37–45 (2014).

67. Datsenko, K. A. & Wanner, B. L. One-step inactivation of chromosomal genes in *Escherichia coli* K-12 using PCR products. Proc Natl Acad Sci U S A 97, 6640–6645 (2000).

68. Molina-García, L. & Giraldo, R. Aggregation interplay between variants of the RepA-WH1 prionoid in *Escherichia coli*. J Bacteriol 196, 2536–2542 (2014).

69. Lobato-Márquez, D., Moreno-Córdoba, I., Figueroa, V., Diáz-Orejas, R. & Garciá-Del Portillo, F. Distinct type I and type II toxin-antitoxin modules control *Salmonella* lifestyle inside eukaryotic cells. Sci Rep 5:1 5, 1–10 (2015).

70. Martell, J. D., Deerinck, T. J., Lam, S. S., Ellisman, M. H. & Ting, A. Y. Electron microscopy using the genetically encoded APEX2 tag in cultured mammalian cells. Nat Protoc 12, 1792–1816 (2017).

71. Salema, V. et al. Selection of single domain antibodies from immune libraries displayed on the surface of E. coli cells with two β-domains of opposite topologies. PLoS One 8, e75126 (2013).

72. López-Jiménez, A. T. et al. High-content high-resolution microscopy and deep learning assisted analysis reveals host and bacterial heterogeneity during *Shigella* infection. Elife 13:RP97495 (2024).

73. Hung, V. et al. Spatially resolved proteomic mapping in living cells with the engineered peroxidase APEX2. Nat Protoc 11:3 11, 456–475 (2016).

74. Olivella, R. et al. QCloud2: an improved cloud-based quality-control system for mass-spectrometry-based proteomics laboratories. J Proteome Res 20, 2010–2013 (2021).

75. Tyanova, S., Temu, T. & Cox, J. The MaxQuant computational platform for mass spectrometry-based shotgun proteomics. Nat Protoc 11, 2301–2319 (2016).

76. Ritchie, M. E. et al. limma powers differential expression analyses for RNA-sequencing and microarray studies. Nucleic Acids Res 43, e47 (2015).

77. Carbon, S. et al. AmiGO: online access to ontology and annotation data. Bioinformatics 25, 288–289 (2009).

78. Perez-Riverol, Y. et al. The PRIDE database at 20 years: 2025 update. Nucleic Acids Res 53, D543 (2024).

79. Fromont-Racine, M., Rain, J. C. & Legrain, P. Toward a functional analysis of the yeast genome through exhaustive two-hybrid screens. Nat Genet 16, 277–282 (1997).

80. Witting, K. F. et al. Generation of the UFM1 Toolkit for Profiling UFM1-Specific Proteases and Ligases. Angew Chem Int Ed Engl 57, 14164–14168 (2018).

81. Hall, C., Flores, M., Storm, T., Crosier, K. & Crosier, P. The zebrafish lysozyme C promoter drives myeloid-specific expression in transgenic fish. BMC Dev Biol 7, 1–17 (2007).

82. Wu, R. S. et al. A rapid method for directed gene knockout for screening in G0 zebrafish. Dev Cell 46, 112–125.e4 (2018).

83. Reinhardt, J. & Kolbe, M. Secretion assay in *Shigella flexneri*. Bio Protoc 4(22): e1302 (2014).

84. Mostowy, S. et al. The zebrafish as a new model for the in vivo study of *Shigella flexneri* interaction with phagocytes and bacterial autophagy. PLoS Pathog 9, e1003588 (2013).

85. Valdivia, R. H. & Falkow, S. Bacterial genetics by flow cytometry: Rapid isolation of *Salmonella* Typhimurium acid-inducible promoters by differential fluorescence induction. Mol Microbiol 22, 367–378 (1996).

86. Drecktrah, D. et al. Dynamic behavior of *Salmonella*-induced membrane tubules in epithelial cells. Traffic 9, 2117–2129 (2008).

87. Thery, F. et al. Ring finger protein 213 assembles into a sensor for ISGylated proteins with antimicrobial activity. Nat Commun 12(1):5772 (2021).

