## Supplementary figures and images for "Proximity biotinylation at the host-*Shigella* interface reveals UFMylation as an antibacterial pathway"

### Extended Data Figures

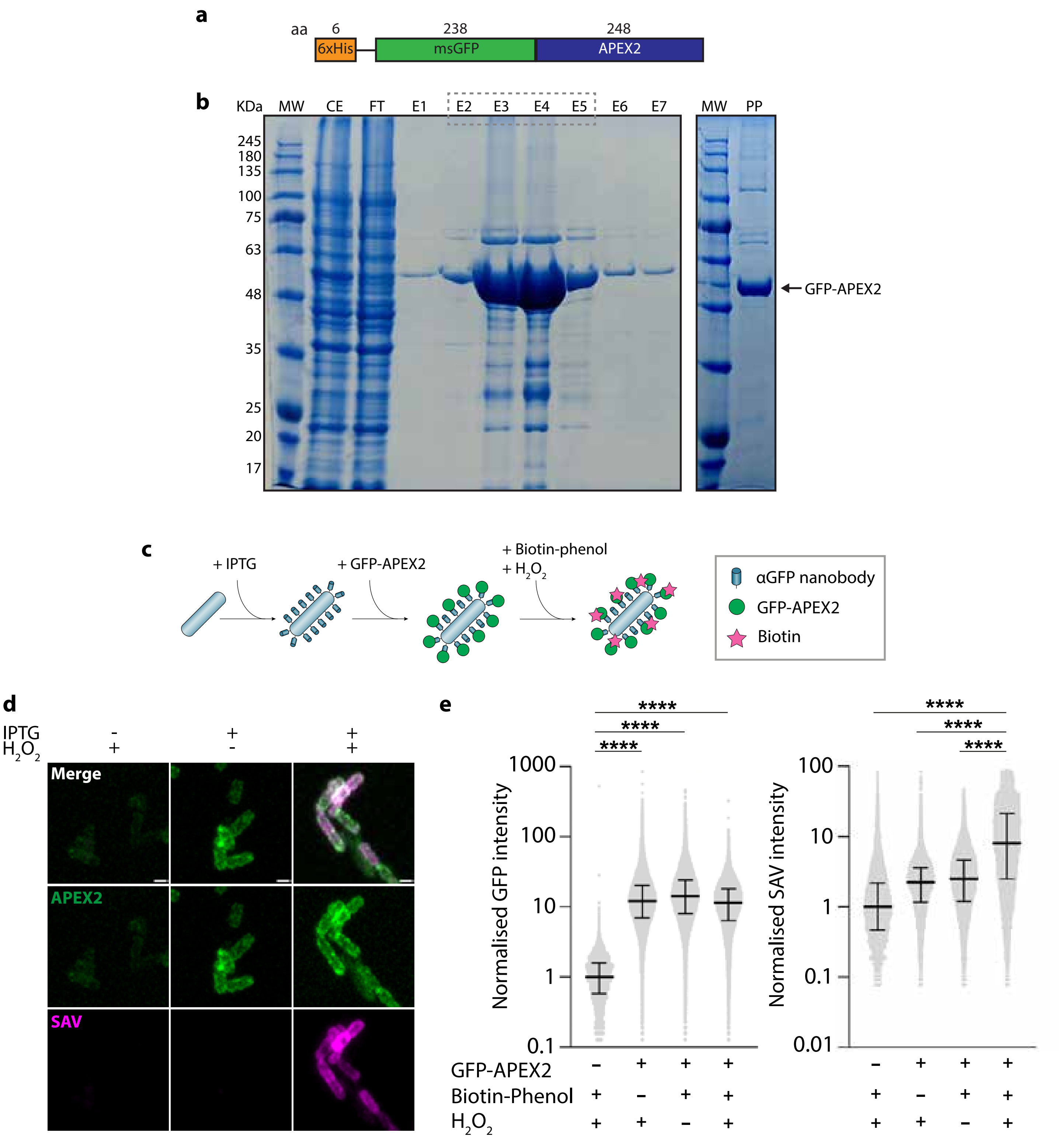

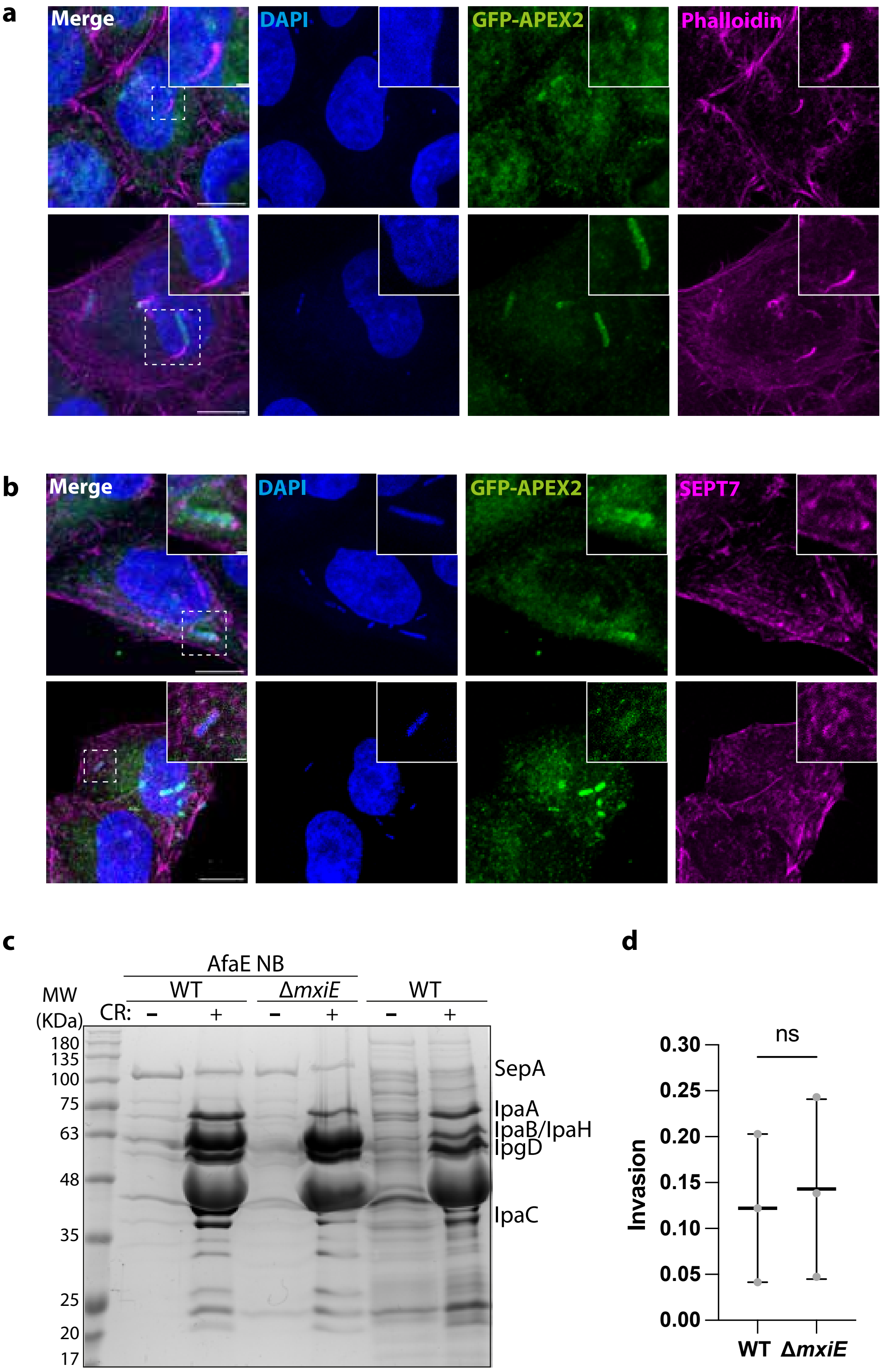

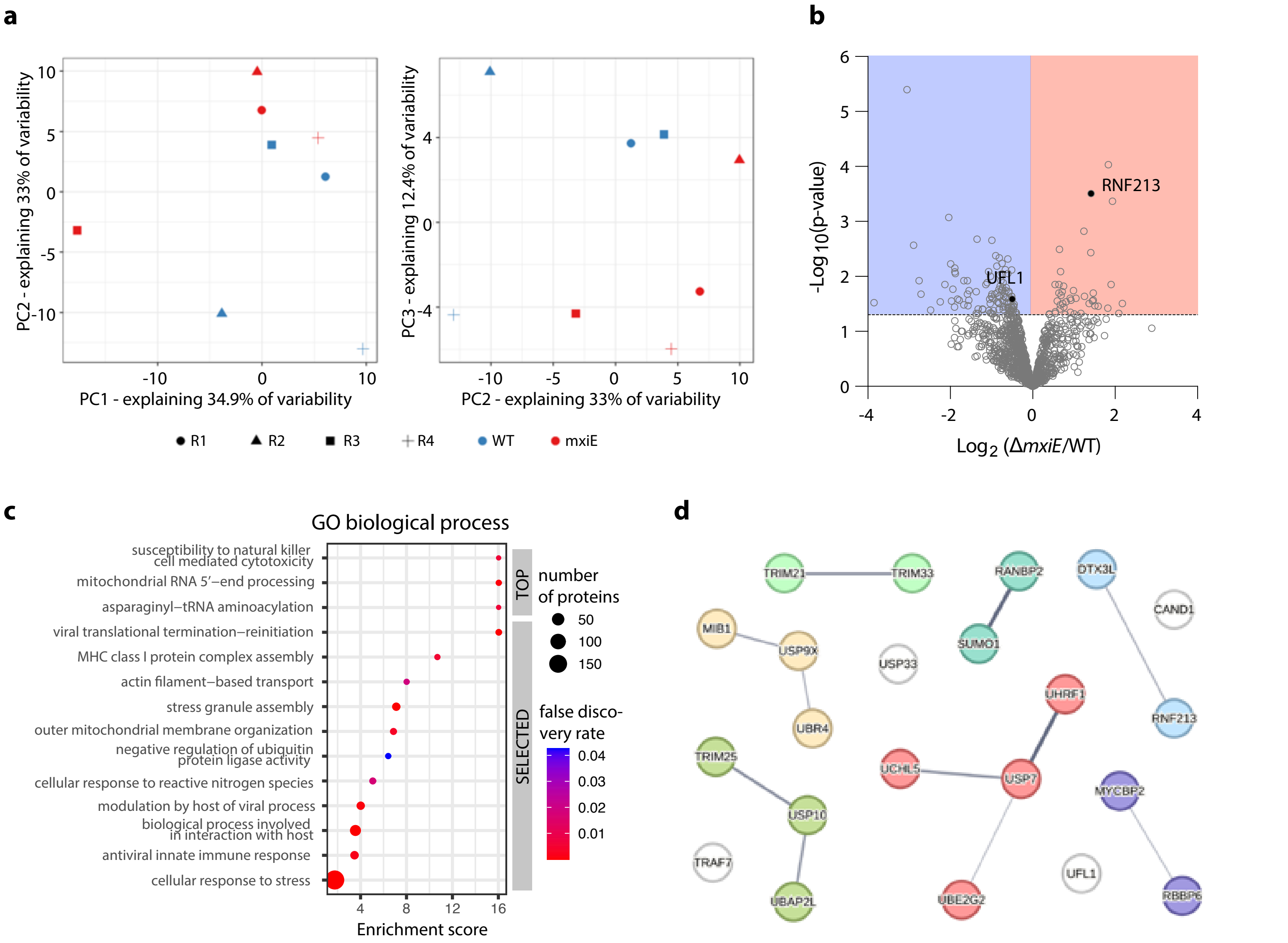

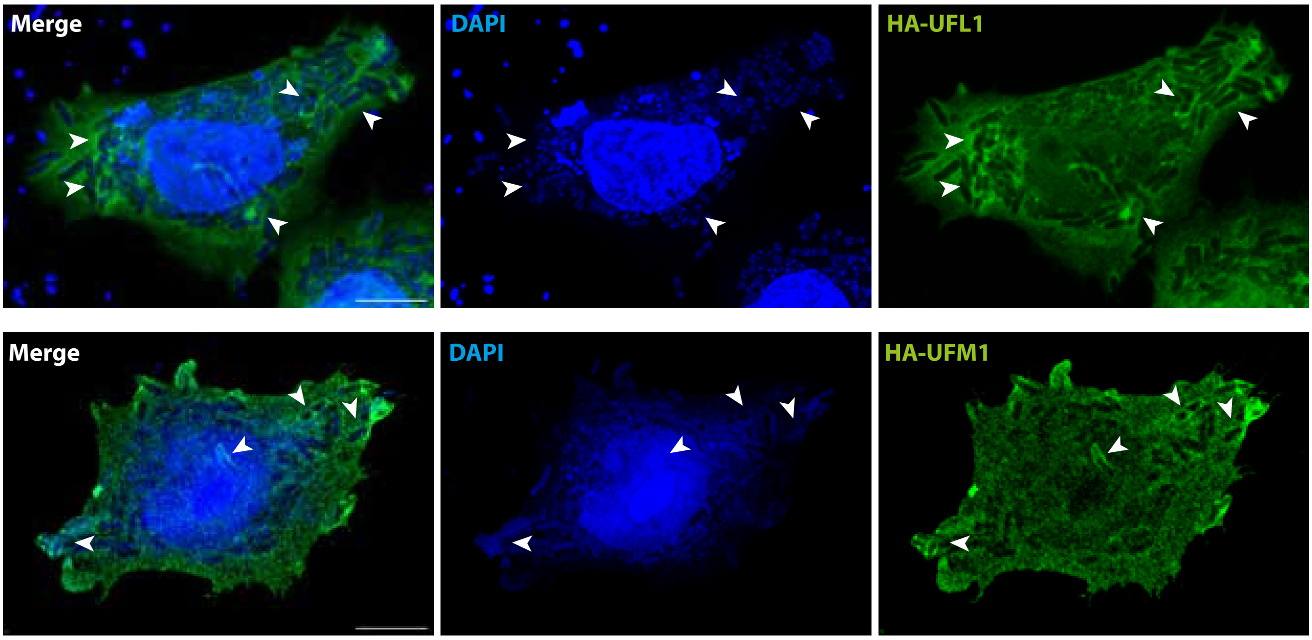

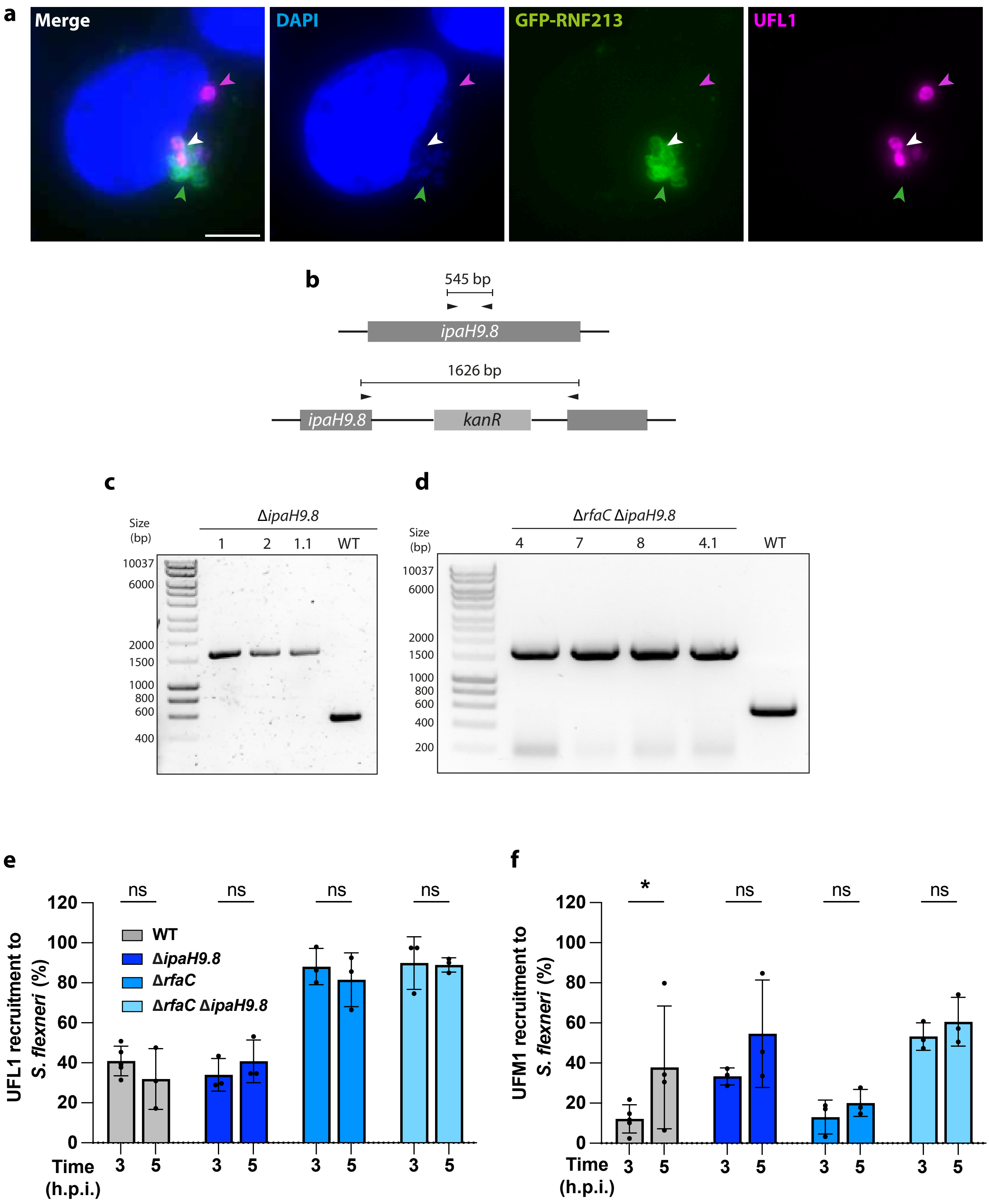

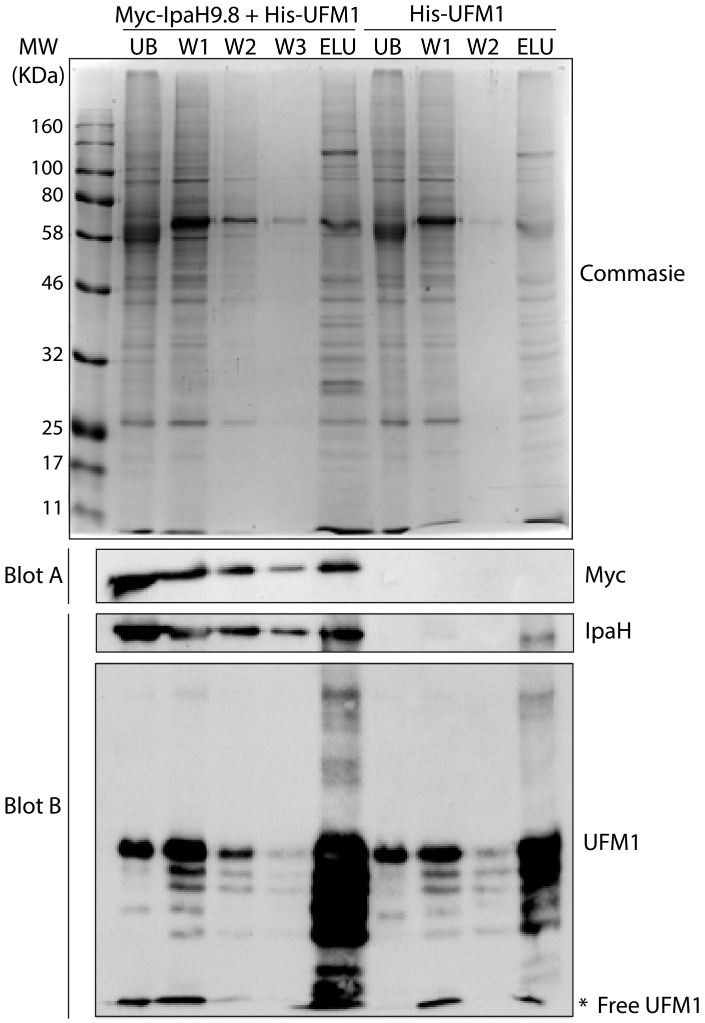

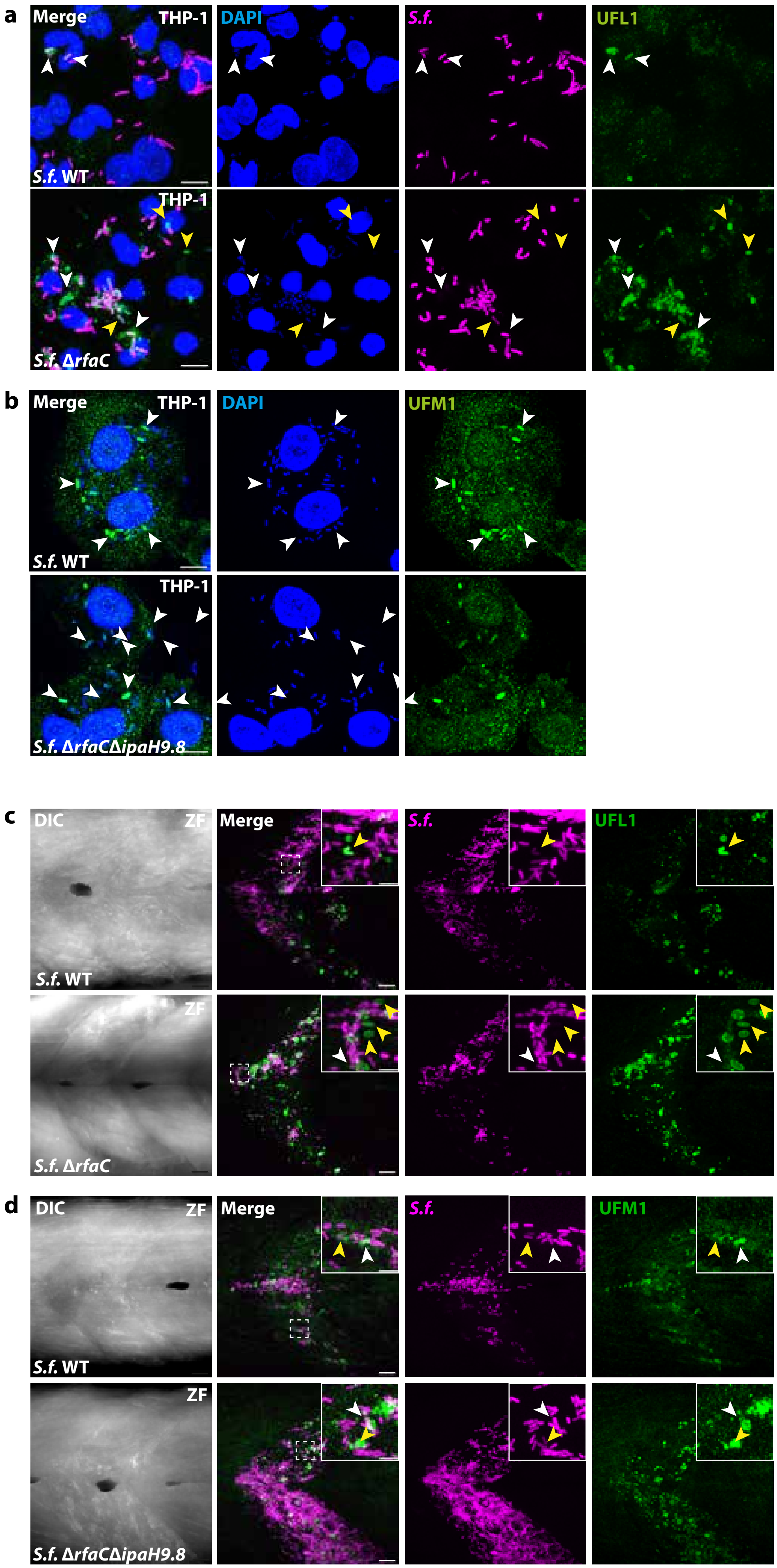

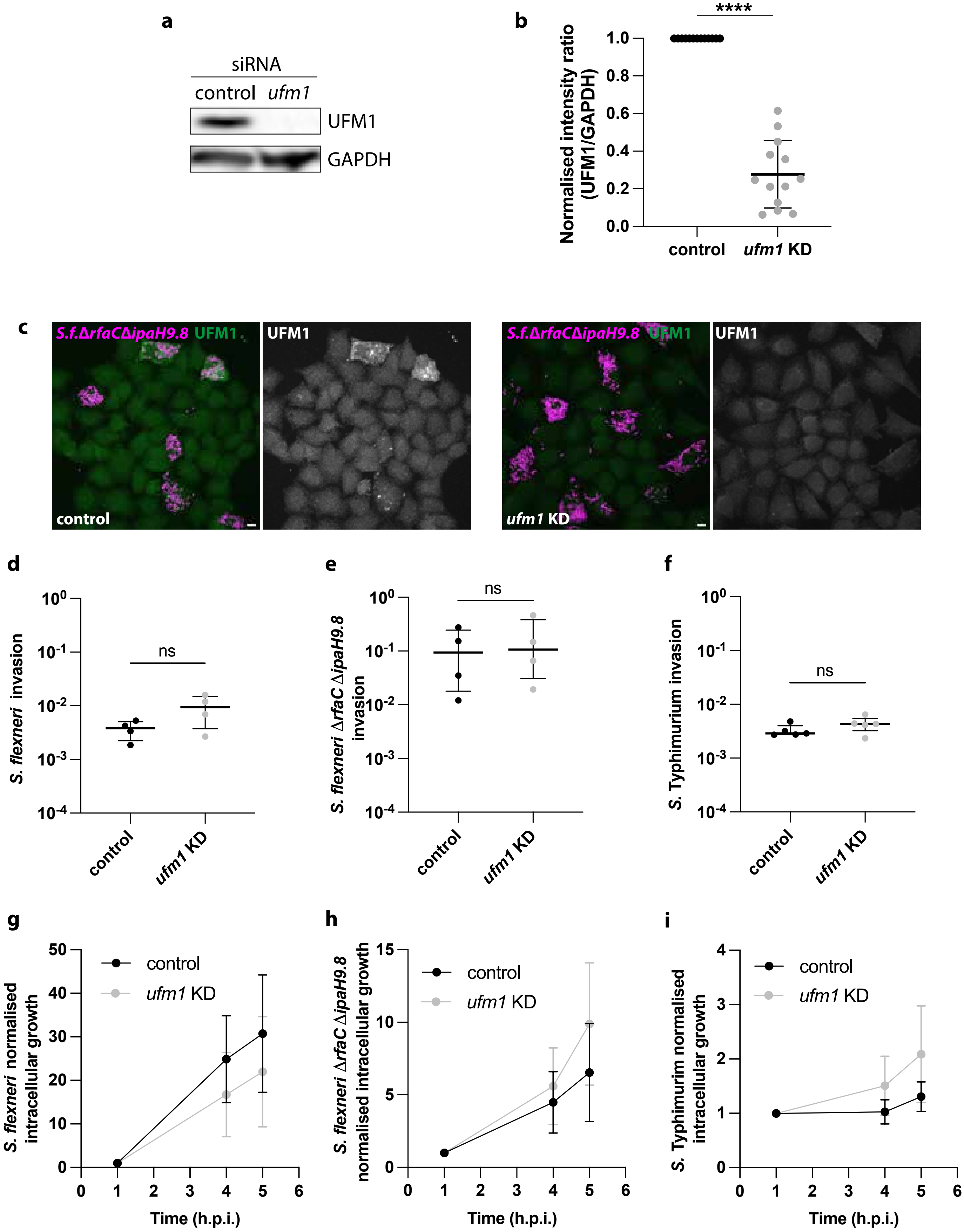

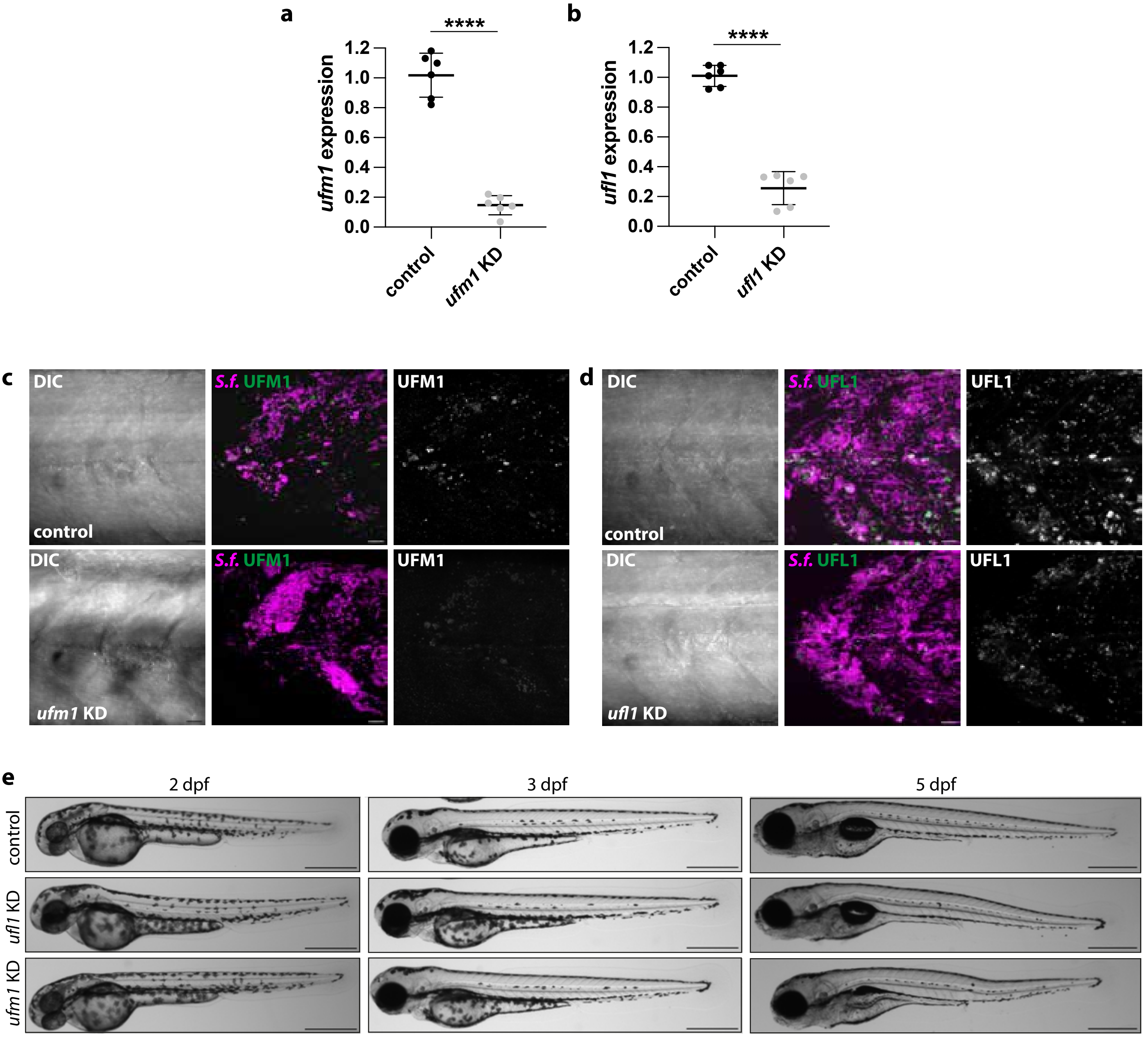

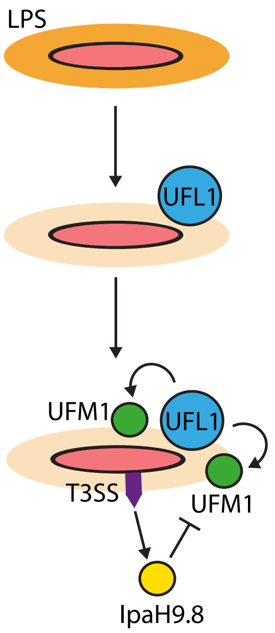
